# Bio-BLIP: A Multimodal Architecture for Transferable Reasoning in Genomic Variant Interpretation

**DOI:** 10.64898/2026.05.12.724740

**Authors:** Anvita Gupta, Alejandro Buendia, Anshul Kundaje, Jure Leskovec

## Abstract

Developing scientific hypotheses in biology requires integrating heterogeneous evidence across DNA sequence, gene context, protein function, and prior literature. Existing multimodal AI systems expose biological evidence to reasoning models through textification or by projecting biological embeddings into fine-tuned language models. However, these models are typically highly optimized the specific set of tasks for which they are fine-tuned. Here we present Bio-BLIP, a multimodal Q-former based architecture which leverages biological embeddings and a LLM to generalize to complex reasoning tasks without task-specific fine-tuning. The key to Bio-BLIP is a new neural network architecture that integrates four data modalities – DNA, genes, proteins, and text – through a master Qformer model, which integrates the modality-specific information into a fixed-length prefix for the LLM backbone. Bio-BLIP is pretrained on the task of human genetic variant annotation and achieves a 29.8% increase in generating accurate variant features over frontier LLMs. We evaluate Bio-BLIP zero-shot on downstream genomic tasks of variant prioritization and target gene prediction. Bio-BLIP outperforms two alignment-free genomic language models on regulatory variant prioritization for Mendelian disease. Across the target gene prediction task, Bio-BLIP improves accuracy over LLMs by leveraging learned genomic variant knowledge in difficult cases. Our model produces rich, transparent reasoning traces. In biological domains characterized by multiple scales of data and varied downstream tasks, Bio-BLIP offers a step toward natively multimodal, generalizable reasoning.

## 1 Introduction

Biology requires reasoning over complex datasets containing diverse multimodal entities, such as DNA, RNA, genes, proteins, tissues, and cells. For example, drug development programs shortlist disease-related target genes by integrating signals across genome-wide association studies, regulatory sequence contexts, protein-level changes, and gene expression. Biologists seek to understand the effects of genomic variants in individuals, which are differences in an person’s DNA sequence when compared to a standard “reference” genome. Variants may be one or hundreds of base-pairs long and affect human traits through multiple scales of biology. Understanding requires the ability to jointly interpret different biological modalities, which remains a core challenge for AI models in science.

Reasoning on diverse datasets is also important for scientific hypothesis generation. Ultimately, AI models that understand diverse biological entities can serve as scientific collaborators, surfacing non-obvious connections and accelerating discovery. Though true collaboration remains distant, large language models (LLMs) have demonstrated reasoning capabilities over textified biological data [1], [2], [3]. However, textualization discards rich geometric and functional representations of biology encoded by state-of-the-art foundation models (FMs) like AlphaGenome [4], AlphaFold [5], and Evo2 [6].

The goal of multimodal architectures in biology is to natively integrate the reasoning capabilities of LLMs with the rich representations contained in biological FMs. Recent works have made progress: BioReason combines an LLM with Evo2 DNA embeddings for variant effect prediction [7], BioReason-Pro extends this to proteins via ESM embeddings [8]. Several GPT-style works connect single-modality FMs to LLMs through lightweight projections and supervised fine tuning [9, 10, 11, 12, 13]. However, these models are constrained to the tasks on which they are trained. Biological FM embeddings have demonstrated broad utility across biological predictions [14, 6, 15, 16]. Theoretically, the general reasoning capabilities of the LLM should be able to leverage Biological FM embeddings to complete varied tasks. However, few multimodal models have shown transfer without further fine-tuning.

Here we introduce Bio-BLIP (Fig. 1), a novel transformer-based architecture for multimodal reasoning that enables zero-shot transfer across related tasks. Bio-BLIP introduces three key innovations. First, it takes in four input modalities simultaneously – DNA sequence embeddings, gene neighborhood information, protein sequences, and text. We simultaneously train several modality-specific Q-former models to align Biological FM embeddings with shared captions. Second, the architecture introduces a new master Q-former model in Stage 2, which integrates modality-specific tokens through crossattention, producing a fixed-length multimodal prefix for the LLM. Finally, the LLM in Bio-BLIP is never fine-tuned with SFT or RL, which allows us to maintain the full reasoning capability of the LLM backbone. This is because the unique training recipe of Bio-BLIP keeps the LLM frozen and only backpropagates next-token loss to the Q-formers. Together, these mechanisms capture task-specific knowledge encoded in Biological FMs, while leveraging the LLM’s reasoning capabilities.

**Figure 1:**
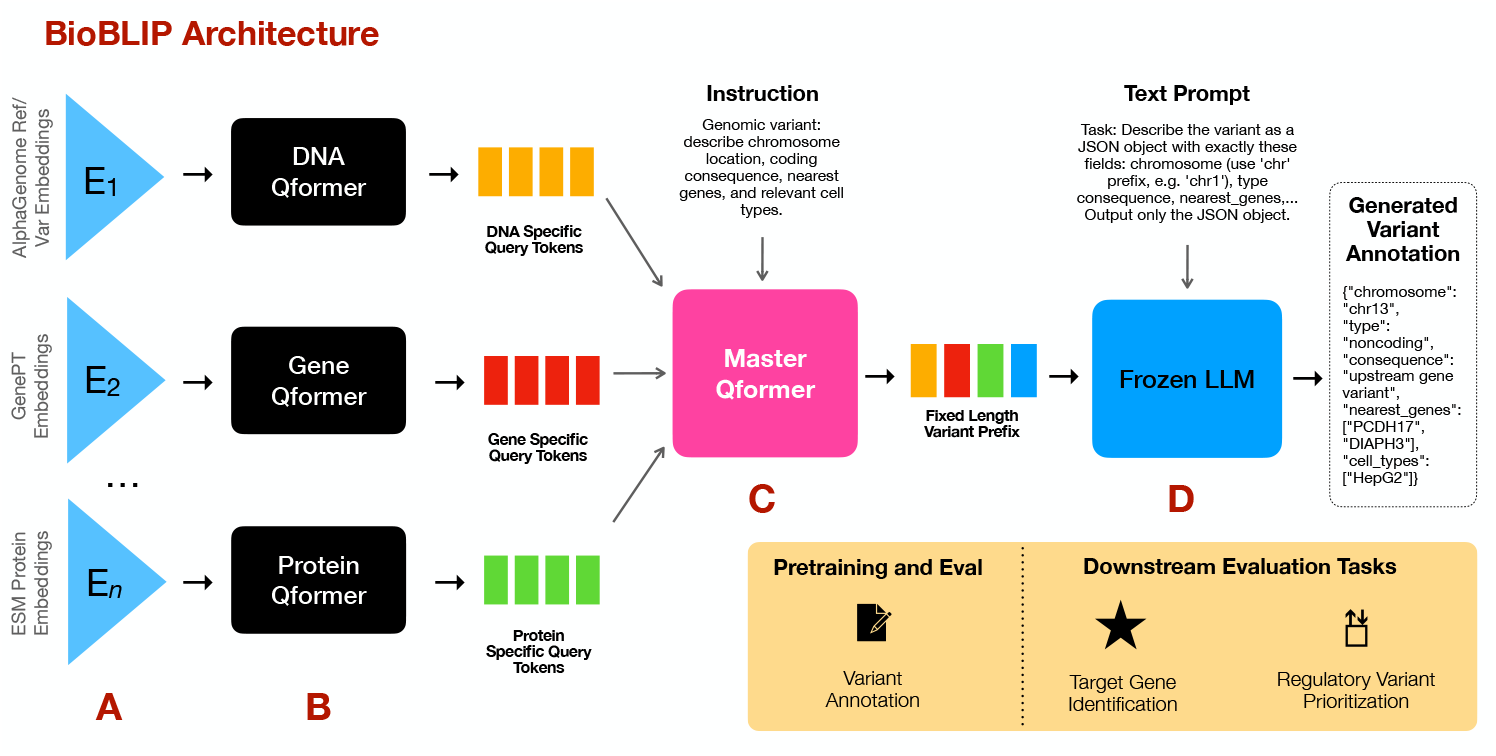
**(a)** Embeddings (E) from Biological Foundation Models (BioFMs) are input to Bio-BLIP to provide multiple modalities of information, here from *AlphaGenome, GenePT*, and *ESM-2*. **(b)** Modality-specific Q-formers align diverse biological embeddings with captions of genomic variants, which contain functional and locational information. The Q-formers output query tokens encoding relevant information from each modality. **(c)** The Master Qformer integrates modality-specific tokens through cross-attention, producing a fixed-length visual prefix **(d)** The LLM, its weights *frozen*, takes as input the multimodal prefix and a prompt. During pretraining, the LLM loss backpropagates only to train the Q-formers. Pretraining and evaluation is conducted on variant annotation; all weights are frozen and the model is then applied to downstream evaluation tasks.

We pretrain Bio-BLIP to annotate single-nucleotide-variants from OpenTargets Genetics and observe strong zero-shot performance on downstream reasoning tasks relating to genomic variants. Bio-BLIP achieves 29.78% improvement across per-field accuracy in variant annotation over Qwen3-32B. For comparison, Claude Sonnet 4.6 achieves comparable performance to Qwen3-32B and 29% lower than Bio-BLIP with equivalent context information expressed as text. Further, Bio-BLIP reduces nearestgene anchoring in OpenTargets Gold Standard target-gene prediction. We also observe that Bio-BLIP outperforms alignment-free genomic language models GPN-Promoter and Evo2-7B on regulatory variant prioritization in the TraitGym Mendelian traits dataset. Biological models that explicitly incorporate genetic conservation remain stronger on some benchmarks, which highlights additional data that can be added to Bio-BLIP’s training. Overall, Bio-BLIP demonstrates a promising new architecture towards building natively multimodal models for reasoning tasks in genomics, moving beyond textification of genomic data and more transferable reasoning.

## 2. Bio-BLIP Model

Bio-BLIP (Fig. 1) is a hierarchical architecture that integrates multiple embeddings of genomic variants for zero shot generalization to neighboring tasks. For this, the training of Bio-BLIP is guided in two stages: **Stage 1** focuses on *modality-specific* text alignment and information extraction, while **Stage 2** integrates modality-specific information through the master model and makes it available to the LLM’s reasoning capabilities.

### 2.1 Pretraining Approach

The space of possible human genomic single-nucleotide variants (SNVs) is combinatorially vast, making exhaustive retrieval or enumeration at inference time difficult. We pretrain Bio-BLIP on variant annotation: a task which allows the model to develop general internal representations of genetic variants. This alignment also promotes generalization. Understanding structural and functional properties of variants is useful for many downstream reasoning tasks with unseen variants.

### 2.2 Pretraining Regime

We train the Bio-BLIP model (Fig. 1) in two stages. These are preceded by an input processing stage which precomputes and stores embeddings from various Biological Foundation Models, here: encoder outputs from AlphaGenome, GenePT-based neighborhood embeddings, and nearest-protein ESM2 embeddings. Precomputing provides significant memory savings, time savings, and greater modularity. This also ensures that we never update the weights of the Biological Foundation Models. Each embedding type receives its own modality-specific *Q-former*.

#### Qformer Definition

A Q-Former (Querying Transformer) is a lightweight transformer module that extracts a fixed-size set of learned representations from variable-length input embeddings. It does so via a set of *K* learnable query vectors 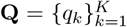, which are randomly initialized and updated during training. These query vectors interact with the input embeddings through cross-attention, effectively “querying” the input for the most task-relevant information. The query vectors also interact with the text caption tokens via cross-attention [17]. The outputs are *modality-specific query tokens*: *K* fixed-dimensional vectors that extract relevant information from the input.

**Stage 1** of training simultaneously trains each modality-specific Q-former model to align its input embeddings with shared textual captions, allowing each to learn modality-specific attention patterns that are relevant to the shared text in different ways. Following the BLIP-2 style of fusing vision and language [17], each Q-former has three losses to optimize in this stage:

- *Image–text contrastive* (ITC) loss maximizes the cosine similarity between each query representation and its paired text caption while minimising similarity to unpaired captions in the batch: 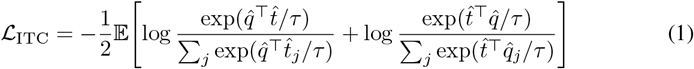 Where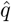 and 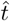 are *l*_2_-normalised query and text projections respectively, and *τ* is a learnable temperature.
- *Image–text matching* (ITM) loss trains the model to identify, given a batch of embeddings and captions, whether a given embedding-caption pair is a match. A binary cross-entropy is applied over positive pairs and negatives mined via ITC similarity scores
- *Image-grounded text generation* (ITG) loss, a causal language-modeling objective in which the text decoder attends to the output query tokens as soft visual prompts and is trained to generate the paired caption autoregressively.

The total Stage 1 loss aggregates all three objectives across each of *N* modality-specific Q-Formers:

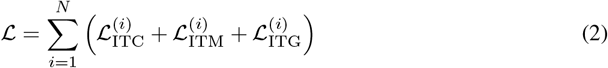

Each Q-former *Q*_*i*_ outputs *K*_*i*_ modality-specific tokens at the end of Stage 1 training.

**Stage 2** of training introduces the Master Q-former model attached to a frozen LLM (Qwen3-4B or Qwen3-32B [18]) for generative variant annotation.

The master Q-former model is a 6-layer InstructBLIP-style [19] Q-Former that fuses the Stage 1 modality-specific tokens with a natural-language instruction, and produces 32 refined query tokens for the LLM. It contains *N*_*m*_ = 32 learnable meta-query vectors of its own. At each forward pass the meta-query tokens and the tokenised instruction are concatenated into a single sequence; the query tokens cross-attend with both the instruction tokens and self-attend with each other (mimicking the InstructBLIP concatenation trick), and cross-attend to the layer-normalised Stage 1 visual tokens every two transformer layers.

The Stage 2 training loss is next-token-prediction with the LLM’s weights frozen, so the loss only updates the projection weights, *W*_proj_, the Master Qformer weights, and the Stage 1 Qformer weights. Notably, the LLM is not trained or fine-tuned during any point in the training regime. We re-use the LLM’s reasoning capability as is. This leads to greater generalization capabilities and is a key difference over existing multimodal models.

#### Input Representations

While the architecture is general and supports any number of input modalities, here we describe the specific inputs to Bio-BLIP:

- **DNA Sequence encoder**. Pre-computed AlphaGenome [4] embeddings for the reference and alternate alleles of a genomic variant, each represented as 256 context-window bins of hidden dimension 3072 and interleaved to form a sequence of length 512.
- **Gene-neighborhood embeddings**. GenePT [20] embeddings for the up to 15 protein-coding genes nearest to the variant. Each embedding is concatenated with log-distance of the gene from the variant and a boolean column which encodes whether the gene physically overlaps the variant or not.
- **Protein-sequence encoder**. ESM-2 [21] embeddings for the reference and alternate aminoacid sequences of the nearest protein-coding gene to the genomic variant.
- **Textual variant captions**. Structured JSON objects for each single nucleotide variant containing chromosome, variant type `(coding/noncoding)`, most severe consequence (e.g. intron variant, missense variant), nearest protein-coding genes, and associated cell types.

#### Training Details

Modality-specific Q-Formers are 12-layer transformers with hidden dimension *d*_*q*_ = 128, 8 attention heads, a feed-forward width of 4*d*_*q*_, and cross-attention to the encoder every two layers. The Alphagenome Q-former has 32 query vectors, while the GenePT Q-former and ESM Q-former are trained with 8 query vectors each. Varying number of query vectors was not found to provide a significant difference in downstream performance. The losses of the modality-specific Qformers are summed and optimised jointly with AdamW with learning rate lr = 10^*−*4^ and weight decay 0.05, under a cosine schedule with a linear warm-up over 10% of total training steps. Stage 1 training proceeds for 15 epochs and requires approximately 4 hours/epoch with two Nvidia A100 GPUs. Stage two training uses AdamW (lr = 10^*−*4^, weight decay 0.05) with a cosine schedule, 10% linear warm-up, gradient clipping at norm 1.0, and mixed-precision (BF16) AMP. The supervision signal is NTP loss on the answer tokens only; prompt tokens are masked with *−* 100. Training proceeds for 15 epochs and requires approximately 3.5 hours/epoch with four Nvidia A100 GPUs.

## 3. Experiments

We complete experiments for Bio-BLIP on three genomic tasks. Variant Annotation (Sec. 3.1) is a pretraining *and* evaluation task. Target gene identification (Sec. 3.2) and variant prioritization (Sec. 3.3) are strictly for zero-shot evaluation.

### 3.1 Pretraining and Evaluation on Variant Annotation

#### Task Definition

Given a variant ID and/or biological embeddings, the model must produce a structured JSON caption for the variant. Ground-truth labels come from the OpenTargets Genetics portal [22]. Each caption field corresponds to a biologically meaningful prediction:

1. **Chromosome identification**. Predict which of the 23 human chromosomes the variant resides on. This is a fundamental localisation task.
2. **Variant type** (coding vs. non-coding). Classify whether the variant is within a proteincoding region or not. Coding variants may alter protein sequence, while non-coding may affect gene regulation.
3. **Most severe consequence**. Predict the most deleterious annotated consequence of the variant (e.g. missense, splice-acceptor, stop-gained). Consequence severity is a primary signal used in clinical variant prioritisation and reflects the likely mechanism of pathogenicity.
4. **Nearest genes**. Identify the genes in proximity to the variant. For non-coding variants in particular, nearest genes are a key indicator of which genes may be disregulated.
5. **Relevant cell types**. Predict which cell types are most likely to be functionally affected by the variant, particularly important in regulatory contexts.

#### Note: Chromosome prediction

Variant IDs are underscore-separated strings of a variant’s chromosome location, position, reference allele, and alternate allele (`chrom_pos_ref_alt`). When variant ID is explicitly provided, an LLM can trivially read the chromosome directly. This is in contrast to Bio-BLIP and certain BioReason models where the Variant ID is not explicitly provided, and chromosome features must be learned from the input embeddings alone. We exclude chromosome from overall accuracy comparisons.

#### Dataset

We train the model on a set of 278k single nucleotide variants (SNVs) from the OpenTargets genetics database [22]. The data processing pipeline samples 300k SNVs from OpenTargets, mapping each to a consequence label and extracting reference and mutant DNA sequences. For non-coding variants, we query QTL credible sets and GWAS-QTL colocalisations to identify relevant tissues and cell types, supplemented by L2G gene predictions. Variants without tissue information from OpenTargets are passed to AlphaGenome, which scores each variant for 5564 ATAC-seq, DNASE, CAGE, CHIP, and splice-site tracks in various cell types and returns the top-5 tracks by absolute effect size. We then apply a genome build match to filter out variants where the gene coordinate lookup tool, genelocator, does not agree with the closest gene labeled by OpenTargets [23]. This filtering step leaves 278k/300k SNVs, which we split randomly into 80% training and 20% validation, with approx. 1k variants held out as a test set for model evaluation.

#### Baselines and Comparative Models

Bio-BLIP architecture and pretraining details are provided in Section 2.2. We compare with BioReason using Evo2-7B embeddings and Qwen3-4B, the best performing model described by the authors [7]. We retrain BioReason for Variant Effect Prediction (Bioreason-VEP) using the authors’ provided datasets and training code. BioReason encodes paired reference and alternate DNA sequences using Evo2 and conditions a language model to generate outputs from these embeddings via SFT. We also pretrain BioReason using SFT on the same OpenTargets Genetics dataset on which Bio-BLIP is trained, producing BioReason-OT for a fair architecture comparison. We train two versions of BioReason-OT on the OpenTargets Dataset: one with the Variant ID provided in the prompt (BioReason-OT + VarID) and one version of BioReason-OT without the variant ID provided in the prompt. This allows us to estimate how much BioReason relies on the variant ID being in the prompt versus the Evo2 embeddings themselves.

We evaluate Qwen3-32B and Claude Sonnet 4.6 LLMs on this task, while also providing both with relevant information in the LLM context by providing the top AlphaGenome predictions for each variant with a 131,072 bp window, capturing local regulatory context across chromatin accessibility, transcription, and transcription factor binding signals.

#### Results and Observations

Variant Annotation Results (Table 1) demonstrate that Bio-BLIP exhibits strong performance across fields, demonstrating effective learning of variant features from biological embeddings alone. The BioReason family of models generally show a high reliance on the prompt, as sampling shows that BioReason-OT experiences mode collapse to a few captions when Variant ID is not provided. Baselines including frontier LLM models fail on the nearest gene task, which is surprising, because later experiments show that frontier models have knowledge of gene-to-position mappings. Bio-BLIP shows the most demonstrable gains on nearest genes, boosting accuracy from 4.25% to 67%. Overall, Bio-BLIP achieves 29.8% increase in accuracy in describing functional and structural variant features compared to frontier LLM baselines. These strong pretraining results set the stage for us to evaluate Bio-BLIP and BioReason-OT+VarID for zero-shot generalization to downstream tasks.

**Table 1:**
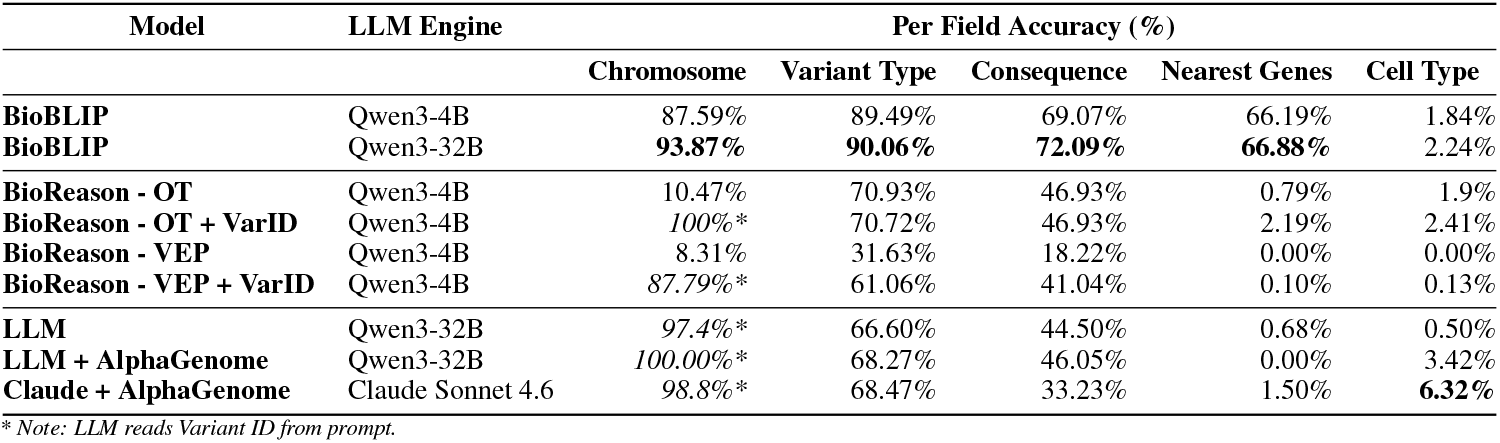
Variant Annotation Results.

Fig. 2 demonstrates that modality-specific Qformers perform better on different aspects of variant annotation. For instance, the Gene Q-former performs best on nearest gene prediction, and AlphaGenome’s Q-former performs best on variant consequence generation. Notably, with the Master Qformer, BioBLIP gains on all of the axes after integrating modality-specific outputs.

**Figure 2:**
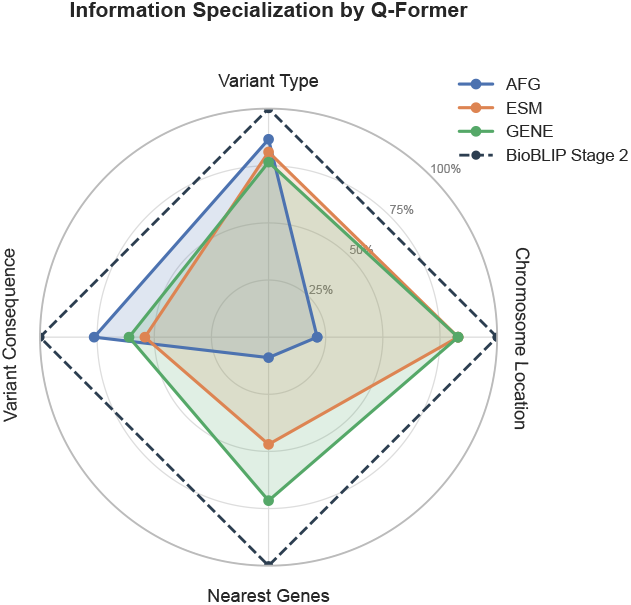
Modality Diversity. AFG = AlphaGenome Embeddings; ESM = ESM-2 Protein Embeddings; GENE = GenePT Neighborhood Embedding. The Qformers for each embedding type show differential per-field accuracy in variant annotation. With the Master Qformer, Bio-BLIP outperforms any specific modality’s model.

**Figure 3:**
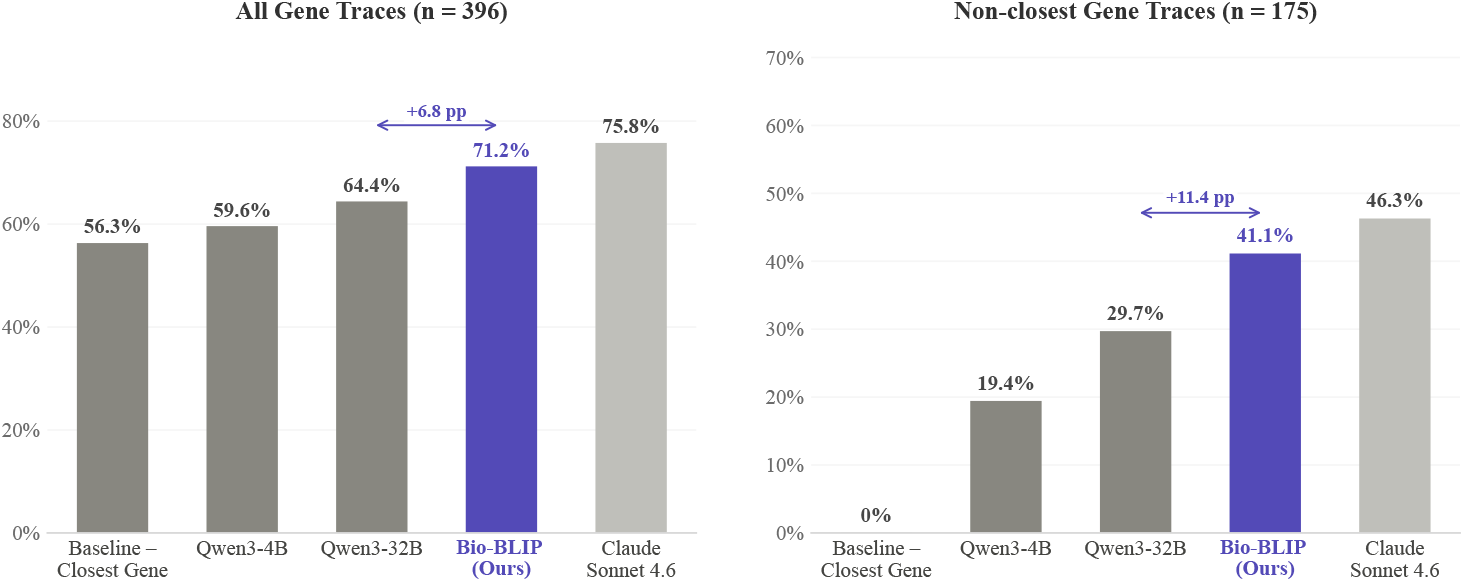
Target Gene Identification. (Left) Accuracy on all reasoning traces. (Right) Accuracy on reasoning traces where the target gene is *not* the closest gene.

### 3.2 Complex Reasoning: Target Gene Identification

#### Task Definition and Dataset

The OpenTargets Genetics Gold Standard dataset contains 397 unique SNVs mapped to their target genes and phenotypes. We construct 5-step chain-of-thought reasoning traces linking GWAS variants from the Gold Standard Dataset to target genes and to the observed phenotype using Claude Sonnet 4.6 and ground-truth expert annotations from the OpenTargets. The task definition is: Given a variant ID, a disease, and a neighborhood of nearby genes, can the model identify the target gene that links the variant to the disease?

#### Baselines

A very strong baseline is simply picking the closest gene by distance to the variant. In 56.31% cases in our dataset, the closest gene is the correct target gene. We also specifically measure performance on traces where the closest gene is not the target gene (“Non-Closest Gene Correct”). Qwen3 baselines and the frontier model Claude Sonnet 4.6 are also tested. Multimodal Baselines include BioReason-OT + VarID, which has been fine-tuned on variant annotation (Sec. 3.1). Multimodal models are also provided the variant and reference allele biological embeddings.

#### Results and Observations

Results for all reasoning traces and non-closest gene traces are shown in Table 2. Notably, Bio-BLIP appears to leverage its multimodal architecture to achieve wins over Qwen3-32B in both overall accuracy (6.8% improvement). It also shows improvement on the more challenging set of non-closest gene traces (11.4% improvement). Bio-BLIP only lags behind the frontier model Claude in a total of nine non-closest gene traces. We used Claude Opus 4.6 to characterize the 23 non-closest gene traces where Bio-BLIP wins over Qwen3-32B, and illustrative traces relating to each pattern are shown in 6.

**Table 2:** Target Gene Identification.

|  | **All Traces** |  | **Non-Closest Gene Correct** |  |
|---|---|---|---|---|
| **Model** | **Accuracy (%)** | **Number Correct** | **Accuracy (%)** | **Number Correct** |
| **Baseline – Closest Gene** | 56.31% | 223/396 | 0.00% | 0/175 |
| **Bio-BLIP Qwen3-32B** | **71.21%** | **282/396** | **41.14%** | **72/175** |
| **BioBLIP Qwen3-4B** | 57.07% | 226/396 | 19.43% | 34/175 |
| **Qwen-32B** | 64.39% | 268/396 | 29.71% | 52/175 |
| **Qwen-4B** | 59.60% | 236/396 | 19.43% | 34/175 |
| **BioReason-OT + VarID** | 2.02% | 8/396 | 0.57% | 1/175 |
| **Claude Sonnet 4.6** | 75.76% | 300/396 | 46.29% | 81/175 |

### 3.3 Regulatory Variant Prioritization

#### Dataset and Task Definition

The TraitGym Mendelian traits benchmark defines a task for regulatory prioritization in human genetics [14]. A Mendelian trait is controlled by a single gene locus and follows simple dominance patterns for inheritance. The benchmark provides 388 groups of candidate regulatory variants with matched location and functions. Exactly one variant per group is labeled “causal” for the given Mendelian trait. The task for the models is to produce a ranked list of all genetic variants from most to least likely to be causal. Evaluation metrics consist of Top-K accuracy and mean rank with the standard deviations provided.

#### Experimental Setup

To extend Bio-BLIP and BioReason for multiple variants, we ran Bio-BLIP once for each variant to compute that variants’ visual prefix and variant summary. The variants’ multimodal prefixes were separated by newline characters and appended to the LLM prompt, as were the variant summaries. BioReason-OT + VarID was also run in the same way, but the multimodal prefixes were replaced by Evo2-7B embeddings for the reference and variant sequences. Variant IDs were provided to all models in the prompts.

#### Baselines

We evaluate BioReason-OT+VarID as sa comparative model, which has been finetuned on the same Opentargets dataset as Bio-BLIP. Biological baselines include Evo2-7B and GPN-Promoter: alignment-free genomic language models. For such models, log-likelihood-ratio (LRR) between variant and the reference sequence is used to predict variant deleteriousness [14]. GPN-MSA is a biological baseline which incorporates evolutionary information [24]. We leverage the precomputed scores for GPN models and Evo2 provided by TraitGym to calculate Top-k accuracies. LLM baselines Qwen3-32B and frontier model Claude Sonnet 4.6 were also tested.

#### Results and Observations

Results (Table 3, Fig. 4) demonstrate that Bio-BLIP shows notably strong zero-shot performance, outperforming GPN-Promoter on both Top-1 and Top-2 Accuracy. We outperform Evo2-7B and Qwen3-32B across the board. We also observe that Bio-BLIP fails to beat GPN-MSA which explicitly uses evolutionary conservation in its predictions. Claude Sonnet performs with incredible 90%+ accuracy on this benchmark, because it immediately identifies the gene linked with a Mendelian trait, finds its coordinates, and ranks variants based on distance. This strategy would not generalize to any trait for which a single causal gene is unknown *a priori*.

**Table 3:** Regulatory Variant Prioritization.

| Model | Top 1 Accuracy | Top 2 Accuracy | Top 3 Accuracy | Mean Rank ± Stdev |
|---|---|---|---|---|
| **Bio-BLIP (Ours)** | ***48.22%*** | ***52.96%*** | *57.40%* | **3.09 ± 2.98** |
| **BioReason-OT + VarID** | 3.55% | 4.44% | 5.03% | 5.15 ± 3.04 |
| **Qwen3-32B** | 19.82% | 28.70% | 36.09% | 4.59 ± 3.06 |
| **Biological Baseline: GPN-Promoter** | 38.76% | 51.18% | **60.06%** | 3.45 ± 2.73 |
| **Biological Baseline: Evo2-7B** | 37.28% | 47.63% | 56.21% | 3.64± 2.76 |
| **Random** | 10.00% | 20.00% | 30.00% | 5.50 ± 2.87 |
| **Biological Baseline: GPN-MSA** | 68.93% | 80.18% | 86.39% | 1.85 ± 1.64 |
| **Claude Sonnet 4.6** | 90.83% | 94.38% | 95.27% | 1.10 ± 0.44 |

**Figure 4:**
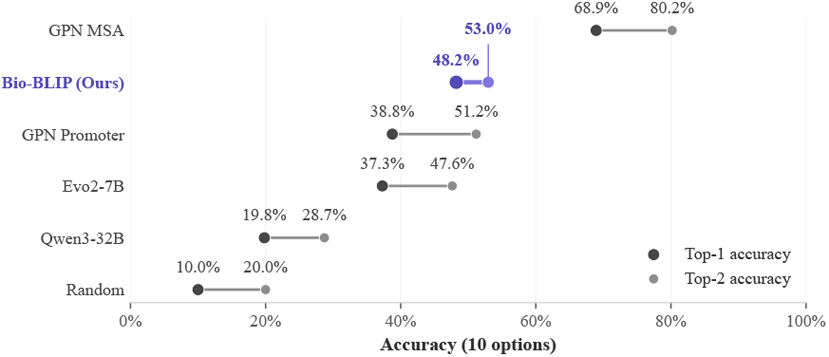
Top-k Accuracy on Regulatory Variant Prioritization in Mendelian Traits.

Bio-BLIP also produces interpretable reasoning traces (illustrative examples provided in App. E.1). Consistently, Bio-BLIP identifies the nearby genes of all variants listed, reasoning about which gene is linked to the Mendelian trait provided, and sometimes reasoning about consequence if that fails. In Trace 2 of the Appendix, where Bio-BLIP identifies two variants putatively close to the same gene HBB for Beta-thalassemia, identifies their consequences (noncoding, intron), and then recalls mutation mechanisms to decide between two HBB-proximal variants.

## 4 Ablation Studies

In this section, we ablate various components of the Bio-BLIP architecture in order to understand the observed performance improvements on benchmark tasks.

### Input Embedding Permutations

We permute the multimodal variant embeddings to investigate how much Bio-BLIP is relying on the biological embeddings as compared to the text prompt. We switch the embeddings away from their respective variants, thus training each annotation on a random variant embedding. As shown in Figure 6 of the Appendix, the ITC loss flatlines. The model is unable to match the true variant annotation with random embeddings.

### Master Q-former Model Ablation

Next, we seek to understand the effect of the Master Qformer model in Bio-BLIP. To probe this, we remove the master model and prepend the tokens from each modality-specific Qformer to the LLM prompt. Results (Fig. 5 and Table 4) demonstrate that performance gains from the master model start with Epoch 2 and lead to an overall 20% increase in performance by the end of the training regime.

**Table 4:** Ablation of Master Qformer Model in Bio-BLIP.

| Method | Chromosome | Type | Consequence | Nearest Genes | Cell Type (ROUGE) |
|---|---|---|---|---|---|
| Bio-BLIP | 48.37% | 86.57% | 62.09% | 42.62% | 5.20% |
| Bio-BLIP: No MM | 30.54% | 85.77% | 60.78% | 7.63% | 4.22% |
| Difference | −17.83% | −0.80% | −1.31% | −34.99% | −0.98% |

**Table 5:** Concordance between BioBLIP and Qwen3-32B on non-closest gene identification (*n* = 175). BioBLIP correctly identifies the causal gene in 23 cases where Qwen3-32B fails, while the reverse occurs in only 3 cases.

|  | **Qwen3-32B Correct** | **Qwen3-32B Incorrect** | **Total** |
|---|---|---|---|
| **BioBLIP Correct** | 49 | 23 | 72 (41.1%) |
| **BioBLIP Incorrect** | 3 | 100 | 103 |
| **Total** | 52 (29.7%) | 123 | 175 |

**Table 6:**
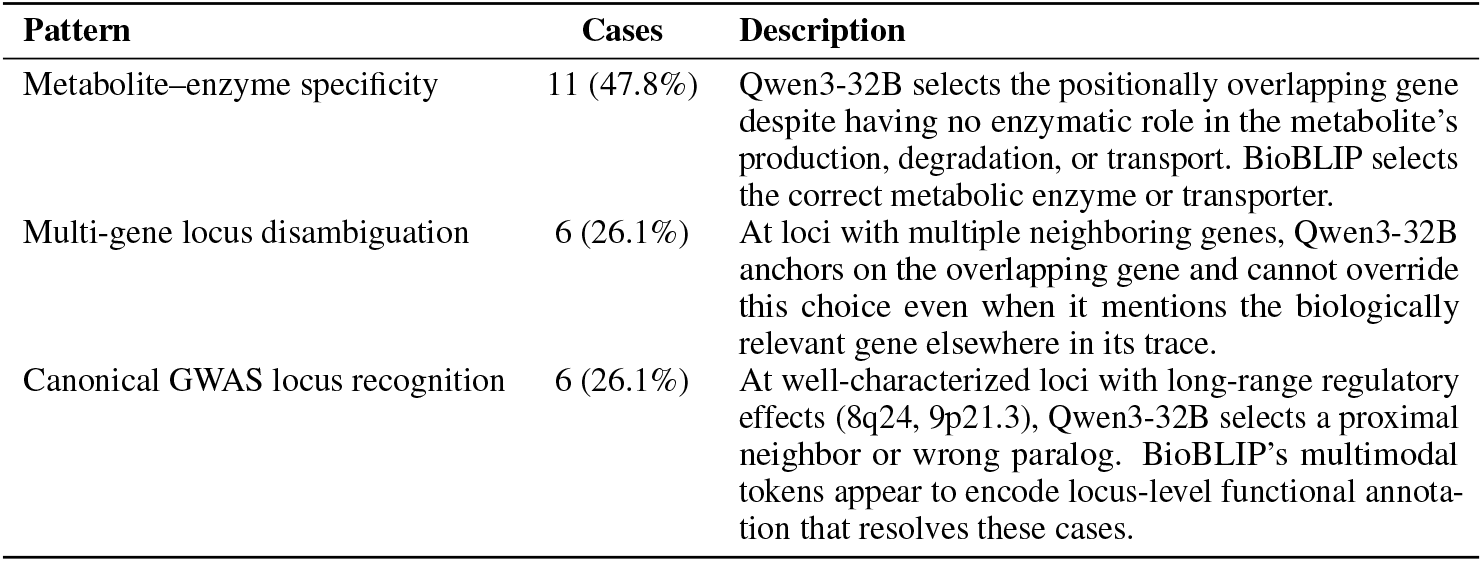
Error pattern breakdown for the 23 cases where BioBLIP identifies the correct causal gene and Qwen3-32B does not. All three patterns stem from Qwen3-32B’s proximity anchoring heuristic.

| **Pattern** | **Cases** | **Description** |
|---|---|---|
| Metabolite–enzyme specificity | 11 (47.8%) | Qwen3-32B selects the positionally overlapping gene despite having no enzymatic role in the metabolite’s production, degradation, or transport. BioBLIP selects the correct metabolic enzyme or transporter. |
| Multi-gene locus disambiguation | 6 (26.1%) | At loci with multiple neighboring genes, Qwen3-32B anchors on the overlapping gene and cannot override this choice even when it mentions the biologically relevant gene elsewhere in its trace. |
| Canonical GWAS locus recognition | 6 (26.1%) | At well-characterized loci with long-range regulatory effects (8q24, 9p21.3), Qwen3-32B selects a proximal neighbor or wrong paralog. BioBLIP’s multimodal tokens appear to encode locus-level functional annota-tion that resolves these cases. |

**Figure 5:**
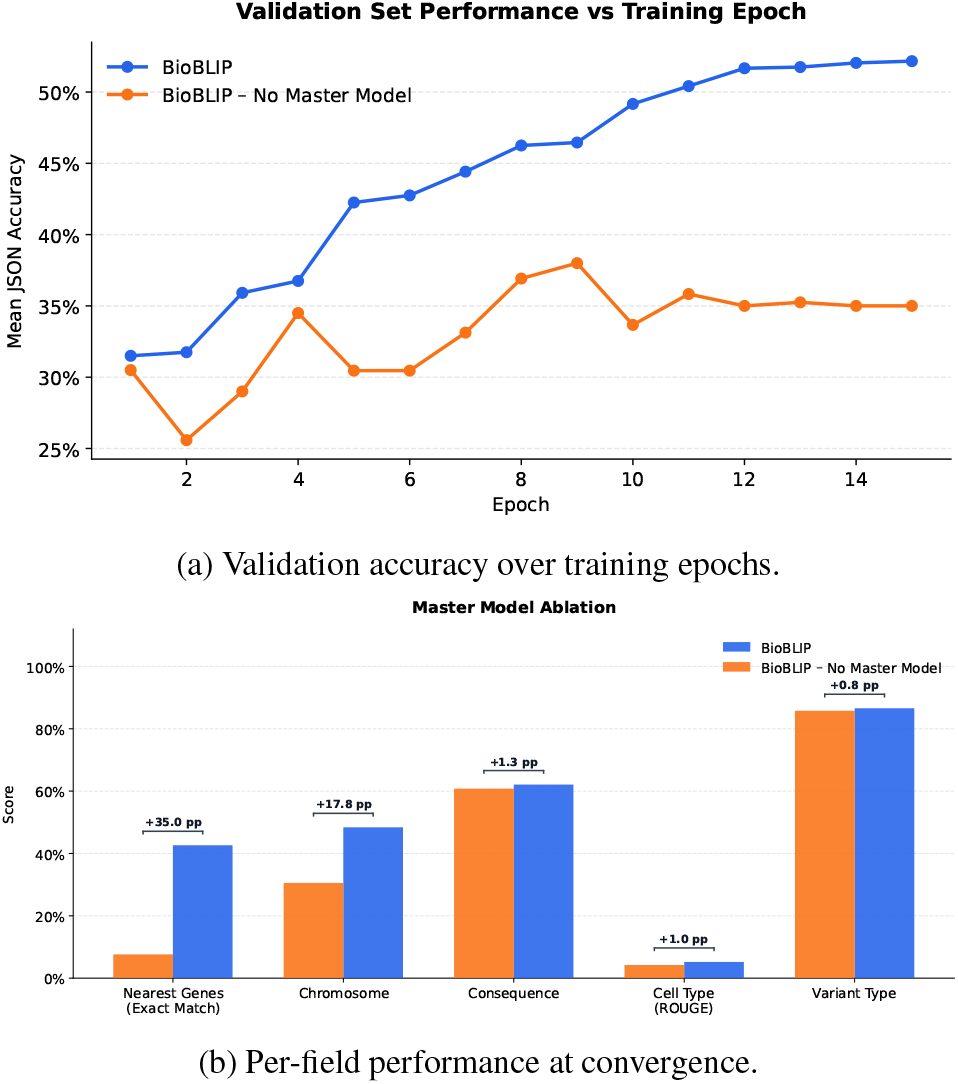
Master Q-former Model ablation. (a) Mean accuracy on the validation set across training epochs for BioBLIP with and without Master Q-former. (b) Per-field breakdown of final performance, with point-improvement (pp) gains. The master model provides substantial gains on fields requiring knowledge from one specific encoder in Stage 1, such as Nearest Genes (+35.0 pp). Fields handled by every encoder (Consequence, Variant Type) demonstrate less difference.

## 5 Related Work

LLaVA-style encoder-projection-LLM architectures [25] have been adapted for biological modalities. BioReason [7] and BioReason-Pro [8] project DNA and protein embeddings through linear layers into an LLM trained with SFT and GRPO, but linear projections provide no mechanism to actively distill task-relevant information from biological embeddings. This incentivizes such models to rely on textual prompt cues and impairs LLM reasoning. [9], [10], [11], and [12] connect biological encoders to LLMs through lightweight projections, but do not evaluate zero-shot transfer to reasoning.

ChatNT [13] fine-tunes the DNA encoder jointly with the projection layer, but does not assess whether pretrained biological representations contribute to performance or whether the system learns task-specific features end-to-end. TxGemma [1], BioT5 [2], and Galactica [3] operate on text-serialized biological inputs, but discards the rich geometric and functional representations learned by specialized biological FMs. Agentic systems such as GeneGPT [26] equip LLMs with database retrieval at inference time, but can only surface information already in existing knowledge bases.

## 6 Discussion and Conclusion

The Bio-BLIP architecture integrates multiple encodings of DNA, gene, and proteins from biological foundation models, exposing these natively to the LLM for transferable reasoning about genomic variants. After pretraining on variant annotation, Bio-BLIP outperforms two alignment-free genomic language models in the Traitgym benchmark of Mendelian traits. We also improve upon Qwen3-32B in identifying distant variant to target gene links, and produce rich multi-step reasoning traces.

While Bio-BLIP is a useful model by itself, limitations exist. Here we evaluated Bio-BLIP on limited benchmarks, but most diseases are complex, influenced by many genes, and not characterized well. Benchmarks should be constructed that are robust and require reasoning about sequence and regulatory context to make correct predictions. The model would need further extensions to natively handle more genomic variants. Interpretability and uncertainty quantification are future directions.

Across tasks and benchmarks, Bio-BLIP demonstrates strong zero-shot performance on neighboring tasks. Due to Bio-BLIP’s novel architecture and training regime, Bio-BLIP maintains full reasoning capabilities of its LLM. Consequently, Bio-BLIP provides a novel transformer-based architecture for genomic variant interpretation that integrates several types of biological embedding information. These advances demonstrate the ability to combine biological foundation models with reasoning capabilities of LLMs, towards transferable reasoning in genomic domains.

## Acknowledgments and Disclosure of Funding

We thank Salil Deshpande, Marcel Roed, Rishabh Ranjan, Vignesh Kothapalli, Sriya Mantena, Moritz Schaefer, Michael Bereket, Rok Sosic, Yanay Rosen, Kuan Pang, Fang Wu, and Aneesh Pappu for discussions and for providing feedback on our manuscript. We also gratefully acknowledge the support of NSF under Nos. CCF-1918940 (Expeditions), DMS-2327709 (IHBEM), IIS-2403318 (III); NIH under No. 1U24NS146314-01, Stanford Data Applications Initiative, Wu Tsai Neurosciences Institute, Stanford Institute for Human-Centered AI, Chan Zuckerberg Initiative, Amazon, Genentech, SAP, and SCBX. A.G. gratefully acknowledges the support of the Knight-Hennessy Fellowship.

The content is solely the responsibility of the authors and does not necessarily represent the official views of the funding entities.

## A Extended Related Work

The encoder-projection-LLM paradigm [25] has seen wide adoption in computational biology. BioReason [7] projects Evo2 DNA embeddings through a linear layer into an LLM which is trained with SFT and GPRO for variant effect prediction, and BioReason-Pro [8] extends this pattern to ESM protein embeddings for protein function reasoning. GPT-style models such as ProteinGPT [9], Prot2Text [10], BioVERSE [11], and MAMMAL [12] similarly connect biological encoders to language models through lightweight linear projections and fine-tune the language model with next-token-prediction. These architectures share fundamental limitations. The linear projection layer provides no mechanism to actively distill task-relevant information from the biological embedding(s). Combined with supervised fine-tuning or LoRA on the LLM backbone, this creates an incentive for the model to rely on textual cues in the prompt and ignore the embeddings. In vision-language models, MIRAGE recently demonstrated that LLMs can attain high scores on multimodal benchmarks in X-ray Q&A even if they completely disregard the image inputs [27]. More systematic ablations of embeddings should be conducted in these Llava-style biological LLMs. These models also do not evaluate zero-shot transfer to reasoning tasks beyond their training tasks.

MAMMAL integrates several biological modalities, such as small molecules, proteins, and cellular representations for regression and classification tasks, but has never been evaluated for complex reasoning tasks [12]. MAMMAL also does not support DNA embeddings. [13] uses the encoderprojection-LLM paradigm to train a conversational DNA agent to answer various types of questions related to nucleic acid sequences. However, ChatNT also fine-tunes the DNA encoder jointly with the projection layer, which makes it difficult to assess whether the pretrained biological representations are meaningfully contributing to predictions or whether the system is simply learning task-specific classification features end-to-end.

A growing body of work applies CLIP-style contrastive alignment to pair biological embeddings with natural language representations. CellWhisperer [28, 29] aligns single-cell transcriptomic profiles with text descriptions, enabling natural language queries over gene expression space. In the protein domain, TourSynBio [30] and ProteinCLIP [31] align protein sequence embeddings with text to support retrieval and editing tasks, and BioBridge leverages such alignment for protein annotation [32]. These approaches are effective for cross-modal retrieval: querying biological entities with natural language or predicting text annotations from biological inputs. However, they are architecturally limited in two respects. The shared embedding space supports matching and retrieval but not multi-step causal reasoning. Second, contrastive alignment becomes combinatorially cumbersome as the number of modalities grows, since each modality pair requires its own alignment objective; with *k* this scales as *O*(*k*^2^) pairwise losses, and the alignment process can distort the internal geometry of pretrained biological embeddings that already encode meaningful structure (e.g., protein homology or gene regulatory relationships).

An alternative to natively multimodal architectures is to convert biological entities into text and reason over them with standard LLMs. TxGemma [1] fine-tunes Gemma on textualized therapeutic data (SMILES strings, protein sequences, disease names) for property prediction and conversational reasoning. BioT5 [2] and Galactica [3] similarly operate on text-serialized biological inputs. While these models benefit from the full reasoning capacity of the LLM, textualization discards rich geometric and functional representations learned by specialized biological foundation models. Textification of heterologous biological datasets is inherently lossy. Agentic systems such as GeneGPT [26] equip LLMs with tool-use capabilities to query biological databases at inference time. These systems can only surface information already catalogued in existing knowledge bases and cannot generalize to novel entities or relationships not present in the literature. Both approaches do not address the fundamental challenge of learning cross-modal biological representations that support generalizable downstream reasoning.

## B Embedding Ablations

We permute the variant text captions and embeddings within each batch, essentially assigning each variant a random caption. We observe in Figure 6 that the ITC loss increases instead of decreasing, as the Q-former is no longer able to provide high cosine similarity to variants matched with the wrong captions.

**Figure 6:**
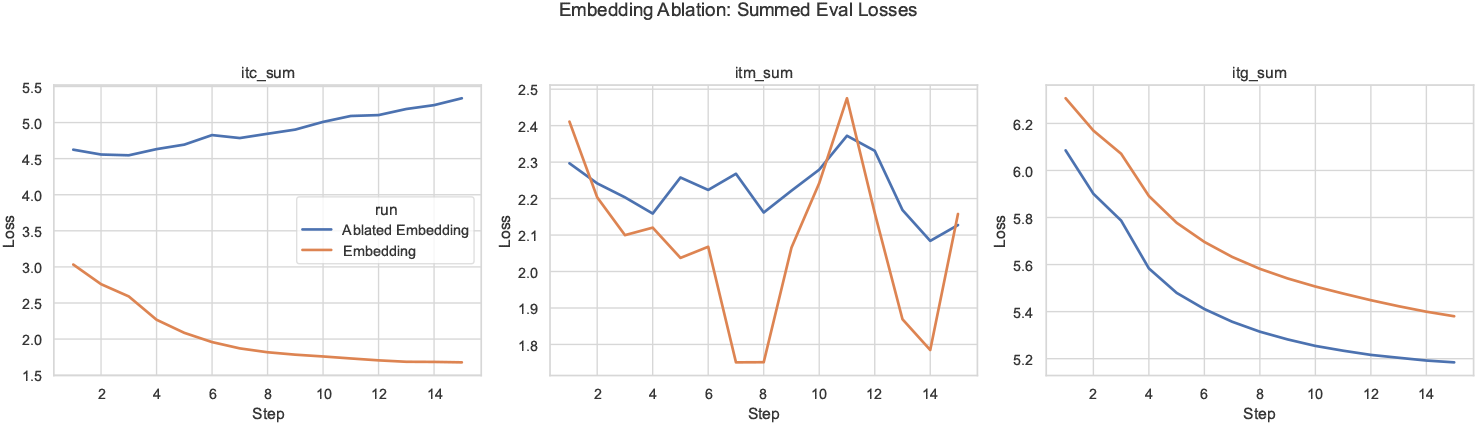
Stage 1 Evaluation Losses for Bio-BLIP with Embeddings permuted to incorrect variants (blue), versus Embeddings matched with the correct variants (orange).

## C Baseline Implementations

### C.1 BioReason SFT Training Details

#### BioReason-VEP

BioReason-VEP was trained with SFT according to the original paper’s VEP-Coding task for classifying coding variants [7]. The input was paired reference and variant DNA embeddings and a textual query providing gene and chromosome context, and the task was to predict if a variant is benign or pathogenic, with its associated disease when pathogenic. Qwen/Qwen3-4B was used for the language model and Evo2-7B-base was used to produce DNA embeddings. BioReason-VEP was trained for 6,106 training steps using DeepSpeed stage 2 and batch size 1. The maximum text length was 1024 tokens and maximum DNA length was 2048 tokens. Evo2 embeddings were extracted from `blocks.28.mlp.l3`. Training was performed with learning rate 5 *×*10^*−*5^, weight decay 0.01, gradient accumulation over 8 steps, LoRA rank 32, LoRA alpha 64, and LoRA dropout 0.05.

#### BioReason-OT

BioReason-OT was trained with SFT on the OpenTargets variant captioning task. The input was paired reference and variant DNA embeddings together with a textual instruction asking the model to describe the variant as a structured JSON object containing chromosome, variant type, consequence, nearest genes, and cell types. The OpenTargets training set was the 250k SNV split augmented with DNA sequence context around the variant. Qwen/Qwen3-4B was used for the language model and Evo2-7B-base was used as the DNA encoder, with Evo2 embeddings extracted from `blocks.28.mlp.l3`. BioReason-OT was trained for 6964 training steps using DeepSpeed stage 2 and batch size 2. The maximum text length was 512 tokens and maximum DNA length was 1024 tokens. Training was performed with learning rate 5 *×*10^*−*5^, weight decay 0.01, gradient accumulation over 8 steps, LoRA rank 32, LoRA alpha 64, and LoRA dropout 0.05.

### C.2 AlphaGenome Prediction Details

AlphaGenome predictions were generated for reference and alternate alleles of each OpenTargets SNV. Each variant was evaluated in a fixed 131,072 bp window centered on the variant and using the hg38 reference genome. Outputs were saved at 128-bp resolution, giving 1024 genomic bins per variant. We included 128-bp resolution bins for ATAC-seq, DNase-seq, CAGE, PRO-cap, RNA-seq, TF ChIP-seq, and histone ChIP-seq tracks. For LLM baselines using AlphaGenome outputs, metadata from the AlphaGenome track metadata table was used to label each output track by assay, biosample, and, when applicable, transcription factor or histone mark [4].

### C.3 LLM-Only Baseline Details

#### Text Only

The text-only baseline used Qwen/Qwen3-32B without DNA embeddings or Al-phaGenome outputs. The model received only the variant identifier string and a JSON-captioning instruction asking for chromosome, coding status, consequence, nearest genes, and cell types. Evaluation used the OpenTargets 1k validation subset. The benchmark was run with maximum generation length 256 tokens.

#### Text Plus AlphaGenome Output Tracks

The AlphaGenome-augmented text baseline used the same Qwen/Qwen3-32B JSON-captioning setup, but the prompt additionally included raw AlphaGenome reference and alternate signal values. For each variant, the top three AlphaGenome tracks were selected by peak absolute log2 fold-change between alternate and reference predictions within a local window of *±* 2 128-bp bins around the variant. The prompt included the raw reference and alternate values for these tracks, labeled with their AlphaGenome assay and biosample metadata, and the model was asked to return only the structured JSON answer. The model was evaluated with maximum generation length 256 tokens.

## D Target Gene Identification Results

### D.1 Illustrative Reasoning Traces

We present six paired reasoning traces produced by BioBLIP and Qwen3-32B for non-closest causal gene identification. Each pair illustrates one of the three failure patterns identified in the Results section. For each case, we provide the variant and trait context, followed by the full five-step trace from each model. The correct prediction is marked with ✓ and the incorrect prediction with *×*.

### D.2 Pattern 1: Metabolite–Enzyme Specificity

#### Case A: Butyrylcarnitine

##### Context

Variant rs2066938 (12_120722812_A_G); trait: butyrylcarnitine; gold standard gene: *ACADS* (acyl-CoA dehydrogenase, short-chain), which catalyses the first step in *β*-oxidation of short-chain fatty acids. Loss of function causes butyrylcarnitine accumulation.

###### BioBLIP (*ACADS* ✓)

*Step 1 — Variant Annotation*. Variant type: noncoding; consequence: intron variant; nearest genes: *ACADS, UNC119B*; cell type: liver hepatocytes.

*Step 2 — Gene-Level Effect*. The variant is located in an intronic region of *ACADS*, a gene encoding a mitochondrial enzyme that catalyses the first step in the *β*-oxidation of short-chain fatty acids, including butyrate. Intronic variants can affect splicing efficiency or regulatory elements, potentially altering *ACADS* expression or protein function. Direction: likely reduced expression or altered splicing of *ACADS*, leading to decreased enzymatic activity.

*Step 3 — Cellular Process*. Impacted pathway: fatty acid *β*-oxidation. Cellular consequence: reduced capacity to metabolise short-chain fatty acids like butyrate, leading to accumulation of butyrylcarnitine and related intermediates in mitochondria.

*Step 4 — Tissue-Level Effect*. Tissue: liver. Outcome: impaired mitochondrial *β*-oxidation in hepatocytes results in altered fatty acid metabolism and increased levels of circulating butyrylcarnitine.

*Step 5 — Disease Mechanism*. The accumulation of butyrylcarnitine due to reduced *ACADS* activity disrupts normal mitochondrial function and energy metabolism in the liver.

Causal gene: *ACADS*.

###### Qwen3-32B (*UNC119B ×*)

*Step 1 — Variant Consequence*. The variant 12_120722812_A_G (rs2066938) is located in the first exon of the *UNC119B* gene, which overlaps with the variant’s genomic position. This is a missense variant. *UNC119B* is the most proximal and overlapping gene, making it the most likely candidate for functional impact.

*Step 2 — Gene-Level Effect. UNC119B* encodes a protein that is a member of the UNC119 family, which is involved in intracellular trafficking and membrane organisation. The missense variant may alter the structure or function of the UNC119B protein, potentially affecting its ability to interact with other proteins or perform its cellular role.

*Step 3 — Cellular Process*. Impacted pathway: intracellular trafficking and membrane organisation. Cellular consequence: disruption of UNC119B function may impair vesicle trafficking or membrane dynamics, which are essential for the proper localisation and function of various membrane-associated proteins.

*Step 4 — Tissue-Level Effect*. Tissue: liver and other metabolically active tissues. Outcome: altered intracellular trafficking in liver cells may disrupt the normal processing and transport of fatty acids or carnitine derivatives, such as butyrylcarnitine.

*Step 5 — Disease Mechanism*. Impaired trafficking or membrane organisation due to UNC119B dysfunction may lead to the accumulation of butyrylcarnitine or its precursors in the liver and bloodstream.

Causal gene: *UNC119B*.

##### Analysis

BioBLIP’s structured annotation identifies the relevant cell type (liver hepatocytes) and connects *ACADS* directly to the *β*-oxidation pathway responsible for butyryl-CoA metabolism. Qwen3-32B notes that the variant “overlaps” *UNC119B*, commits to that gene in Step 1, and constructs a multi-step but biologically implausible pathway from vesicle trafficking to butyrylcarnitine accumulation. *UNC119B* has no established role in fatty acid oxidation or carnitine metabolism.

#### Case B: Isobutyrylcarnitine

##### Context

Variant rs2404602 (15_76738451_A_G); trait: isobutyrylcarnitine; gold standard gene: *ETFA* (electron transfer flavoprotein subunit alpha), which accepts electrons from acyl-CoA dehydrogenases during *β*-oxidation of branched-chain fatty acids. ETFA deficiency causes isobutyrylcarnitine accumulation.

###### BioBLIP (*ETFA* ✓)

*Step 1 — Variant Annotation*. Variant type: noncoding; consequence: intron variant; nearest genes: *ETFA, SCAPER, RCN2, PSTPIP1, TSPAN3, PEAK1, ISL2, TMEM266, NRG4, HMG20A, FBXO22, LINGO1*; cell type: liver.

*Step 2 — Gene-Level Effect*. The variant is an intronic variant in *ETFA* (Electron Transfer Flavoprotein Alpha Subunit), a key component of the electron transfer flavoprotein (ETF) complex. This complex is essential for the transfer of electrons from acyl-CoA dehydrogenases to the mitochondrial respiratory chain, particularly in the oxidation of fatty acids and branched-chain amino acids. Direction: likely reduced expression or impaired function of ETFA.

*Step 3 — Cellular Process*. Impacted pathway: fatty acid oxidation and branched-chain amino acid metabolism. Cellular consequence: reduced activity of the ETF complex leads to impaired electron transfer from acyl-CoA dehydrogenases, resulting in the accumulation of acylcarnitines, including isobutyrylcarnitine, a byproduct of isoleucine metabolism.

*Step 4 — Tissue-Level Effect*. Tissue: liver. Outcome: impaired fatty acid oxidation and branched-chain amino acid metabolism lead to accumulation of isobutyrylcarnitine in plasma.

*Step 5 — Disease Mechanism*. The accumulation of isobutyrylcarnitine reflects a defect in mitochondrial energy metabolism, particularly in the liver.

Causal gene: *ETFA*.

###### Qwen3-32B (*SCAPER ×*)

*Step 1 — Variant Consequence*. The variant 15_76738451_A_G (rs2404602) is located within the gene *SCAPER*, which is annotated as overlapping with the variant. This suggests that the variant is likely a coding variant within the *SCAPER* gene.

*Step 2 — Gene-Level Effect. SCAPER* (Scaffolding protein for AP-1 and ERK) is a scaffold protein that facilitates the interaction between ERK and AP-1, playing a role in the regulation of ERK signalling. A missense variant in *SCAPER* could disrupt its scaffolding function, altering the efficiency of ERK signalling.

*Step 3 — Cellular Process*. Impacted pathway: ERK/MAPK signalling pathway. Cellular consequence: disruption of ERK signalling could lead to altered cellular responses to growth factors and stress signals, potentially affecting metabolic regulation and mitochondrial function.

*Step 4 — Tissue-Level Effect*. Tissue: liver and muscle tissues. Outcome: altered ERK signalling in these tissues may disrupt metabolic homeostasis, particularly in the regulation of fatty acid metabolism and carnitine-dependent pathways.

*Step 5 — Disease Mechanism*. The disruption of ERK signalling due to altered *SCAPER* function may lead to dysregulation of carnitine metabolism, specifically affecting the transport or utilisation of isobutyrylcarnitine.

Causal gene: *SCAPER*.

##### Analysis

Both genes appear in BioBLIP’s candidate list; BioBLIP selects *ETFA* because it encodes a direct component of the enzymatic machinery for isobutyryl-CoA oxidation. Qwen3-32B picks *SCAPER* as the “overlapping” gene and attempts to link a signalling scaffold protein to carnitine metabolism through a long and indirect causal chain (ERK signalling *→* metabolic homeostasis *→* carnitine pathways). *SCAPER* has no established role in acylcarnitine metabolism.

### D.3 Pattern 2: Multi-Gene Locus Disambiguation Case C: LDL Cholesterol (1p13.3 locus)

#### Context

Variant rs646776 (1_109275908_C_T), the canonical 1p13.3 LDL locus; trait: LDL cholesterol; gold standard gene: *SORT1* (sortilin), which controls hepatic VLDL/LDL secretion. The rs12740374 LD block at this locus is one of the best-characterised noncoding causal mechanisms in cardiovascular genetics.

##### BioBLIP (*SORT1* ✓)

*Step 1 — Variant Annotation*. Variant type: noncoding; consequence: intron variant; nearest genes: *CELSR2, PSRC1, MYBPHL, SORT1, SARS, KIAA1324, PSMA5, C1orf194, TMEM167B, SYPL2, ATXN7L2*; cell type: liver hepatocytes.

*Step 2 — Gene-Level Effect*. The variant is located in the intronic region of *CELSR2*, which is part of a gene cluster with *PSRC1* and *SORT1. SORT1* encodes a protein involved in the sorting and secretion of low-density lipoprotein (LDL) particles in the liver. This intronic variant may alter splicing or regulatory elements that influence *SORT1* expression. Direction: likely a gain of regulatory function, increasing *SORT1* expression or altering its splicing.

*Step 3 — Cellular Process*. Impacted pathway: lipoprotein metabolism and secretion. Cellular consequence: increased *SORT1* expression leads to enhanced retention of LDL particles in the liver, reducing their secretion into the bloodstream.

*Step 4 — Tissue-Level Effect*. Tissue: liver. Outcome: reduced secretion of LDL particles into the systemic circulation, resulting in lower plasma LDL cholesterol levels.

*Step 5 — Disease Mechanism*. The intronic variant in *CELSR2* indirectly lowers LDL cholesterol by modulating *SORT1* expression in hepatocytes, thereby reducing LDL secretion.

Causal gene: *SORT1*.

##### Qwen3-32B (*CELSR2 ×*)

*Step 1 — Variant Consequence*. The variant rs646776 is located 157 bp upstream of the *CELSR2* gene, placing it in the promoter or regulatory region. Given its proximity to *CELSR2*, this variant is most likely to influence the expression of *CELSR2*.

*Step 2 — Gene-Level Effect. CELSR2* encodes a member of the cadherin-eptin superfamily, involved in cell adhesion and signalling. It is known to interact with *SORT1*, a gene involved in the trafficking and secretion of LRP1, which regulates LDL cholesterol levels. Direction: the variant is associated with decreased *CELSR2* expression, which in turn reduces the interaction with *SORT1*, leading to increased hepatic secretion of PCSK9.

*Step 3 — Cellular Process*. Impacted pathway: LDL cholesterol metabolism and PCSK9-mediated regulation of LDLR degradation. Cellular consequence: reduced *CELSR2* expression leads to increased PCSK9 levels, promoting LDLR degradation and reducing LDL cholesterol clearance.

*Step 4 — Tissue-Level Effect*. Tissue: liver. Outcome: increased levels of circulating LDL cholesterol due to reduced LDLR activity.

*Step 5 — Disease Mechanism*. Increased LDL cholesterol levels resulting from reduced *CELSR2* expression and elevated PCSK9 activity contribute to dyslipidaemia.

Causal gene: *CELSR2*.

#### Analysis

BioBLIP’s Step 1 lists both *CELSR2* and *SORT1* as candidate genes and annotates liver hepatocytes as the cell type. In Step 2, it explicitly acknowledges that the variant lies “in the intronic region of *CELSR2*” but identifies *SORT1* as the biologically relevant gene for LDL metabolism. Qwen3-32B anchors on *CELSR2* due to positional proximity and fabricates a CELSR2–SORT1– PCSK9 pathway that is biologically implausible. Notably, Qwen3-32B *mentions SORT1* in its trace but cannot override the proximity anchor to select it.

#### Case D: Suntan

##### Context

Variant chr20_34077942_A_G (3^*′*^ UTR variant); trait: suntan; gold standard gene: *ASIP* (Agouti Signalling Protein), an antagonist of MC1R. Increased ASIP expression suppresses eumelanin synthesis and reduces tanning capacity.

###### BioBLIP (*ASIP* ✓)

*Step 1 — Variant Annotation*. Variant type: noncoding; consequence: 3^*′*^ UTR variant; nearest genes: *ASIP, RALY, EIF2S2*; cell type: melanocytes.

*Step 2 — Gene-Level Effect*. The variant is located in the 3^*′*^ UTR of *ASIP* (Agouti Signalling Protein), a key regulator of melanin production. Variants in the 3^*′*^ UTR can affect mRNA stability, translation efficiency, or microRNA binding, potentially altering ASIP expression levels. Direction: likely increased expression of ASIP.

*Step 3 — Cellular Process*. Impacted pathway: melanocortin signalling, specifically through interaction with the MC1R receptor. ASIP acts as an antagonist of MC1R, which normally promotes eumelanin (dark pigment) production. Cellular consequence: increased ASIP expression leads to reduced MC1R signalling, favouring pheomelanin (red/yellow pigment) production over eumelanin in melanocytes.

*Step 4 — Tissue-Level Effect*. Tissue: skin (melanocytes in the epidermis). Outcome: altered melanin type and distribution, resulting in reduced tanning capacity and increased susceptibility to UV-induced damage.

*Step 5 — Disease Mechanism*. Increased ASIP expression reduces the skin’s ability to produce protective eumelanin in response to UV exposure, leading to reduced capacity to develop a suntan.

Causal gene: *ASIP*.

###### Qwen3-32B (*RALY ×*)

*Step 1 — Variant Consequence*. The variant 20_34077942_A_G overlaps with the gene *RALY*, which is an RNA-binding protein involved in post-transcriptional regulation of gene expression. Since it is annotated as overlapping *RALY*, it is most likely a coding or regulatory variant in *RALY*.

*Step 2 — Gene-Level Effect. RALY* is an RNA-binding protein that regulates the stability and translation of specific mRNAs, including those involved in pigmentation and skin biology. A variant in *RALY* could alter its RNA-binding affinity or stability, affecting the regulation of downstream targets.

*Step 3 — Cellular Process*. Impacted pathway: melanogenesis and pigmentation signalling pathways, including regulation of MITF and MC1R expression. Cellular consequence: altered RALY function could lead to dysregulated mRNA stability or translation of key pigmentation genes, resulting in increased or decreased melanin production.

*Step 4 — Tissue-Level Effect*. Tissue: skin (melanocytes). Outcome: changes in melanin synthesis and distribution in the skin, leading to increased or decreased pigmentation in response to UV exposure.

*Step 5 — Disease Mechanism*. Altered regulation of pigmentation-related mRNAs by RALY leads to modulated melanin production in response to UV light, resulting in a genetically determined trait of increased suntan ability.

Causal gene: *RALY*.

##### Analysis

BioBLIP correctly identifies the variant as a 3^*′*^ UTR variant in *ASIP* and the cell type as melanocytes, producing a specific mechanistic chain: ASIP ⊣ MC1R → eumelanin. Qwen3-32B picks *RALY* because it “overlaps” the variant and constructs a generic post-transcriptional regulation story. Qwen3-32B invokes the correct pigmentation pathway (MITF, MC1R) but attributes it to the wrong gene—it possesses the relevant biological knowledge but cannot use it to override the proximity anchor.

### D.4 Pattern 3: Canonical GWAS Locus Recognition

#### Case E: Prostate Carcinoma (8q24 locus)

##### Context

Variant chr8_127310936_T_C; trait: prostate carcinoma; gold standard gene: *MYC*. The 8q24 locus is one of the most-replicated cancer GWAS loci; noncoding variants here act as tissue-specific super-enhancers driving *MYC* overexpression.

###### BioBLIP (*MYC* ✓)

*Step 1 — Variant Annotation*. Variant type: noncoding; consequence: intron variant; nearest genes: *MYC*; cell type: prostate epithelial cells.

*Step 2 — Gene-Level Effect*. The variant is located in an intronic region of *MYC*, which encodes a transcription factor that regulates cell proliferation, apoptosis, and metabolism. Intronic variants can affect splicing efficiency or alter regulatory elements. Direction: likely increased expression or dysregulated splicing of *MYC*, leading to a gain of function.

*Step 3 — Cellular Process*. Impacted pathway: MYC signalling, including regulation of the cell cycle (G1/S transition), ribosome biogenesis, and metabolic reprogramming. Cellular consequence: increased cell proliferation and reduced apoptosis in prostate epithelial cells.

*Step 4 — Tissue-Level Effect*. Tissue: prostate gland epithelium. Outcome: hyperproliferation of epithelial cells, increasing the risk of malignant transformation.

*Step 5 — Disease Mechanism*. Dysregulation of *MYC* expression leads to sustained activation of oncogenic pathways in prostate epithelial cells, promoting clonal expansion and contributing to prostate carcinoma.

Causal gene: *MYC*.

###### Qwen3-32B (*POU5F1B ×*)

*Step 1 — Variant Consequence*. The variant 8_127310936_T_C is located on chromosome 8 at position 127,310,936. It is in close proximity (11,247 bp) to the gene *POU5F1B*, which is a transcription factor involved in developmental processes and cell fate determination. Given the proximity to the transcription start site of *POU5F1B*, this variant is likely a regulatory variant, potentially affecting enhancer or promoter activity.

*Step 2 — Gene-Level Effect. POU5F1B* is a member of the POU domain family of transcription factors, known to regulate genes involved in cell proliferation and differentiation. A regulatory variant in its vicinity could alter its expression levels, potentially leading to dysregulated downstream gene networks. Direction: increased expression of *POU5F1B*.

*Step 3 — Cellular Process*. Impacted pathway: cell proliferation and differentiation, including Wnt/*β*-catenin and Notch signalling. Cellular consequence: increased *POU5F1B* expression may lead to enhanced cell proliferation and reduced differentiation.

*Step 4 — Tissue-Level Effect*. Tissue: prostate epithelial cells. Outcome: dysregulated *POU5F1B* expression may lead to uncontrolled cell growth and reduced apoptosis, contributing to prostatic intraepithelial neoplasia (PIN).

*Step 5 — Disease Mechanism*. Increased *POU5F1B* expression due to the regulatory variant may drive proliferative and anti-apoptotic signalling in prostate epithelial cells, leading to prostate carcinoma.

Causal gene: *POU5F1B*.

##### Analysis

ioBLIP lists only *MYC* as the nearest gene and identifies prostate epithelial cells as the cell type, correctly reflecting the known regulatory architecture at 8q24. Qwen3-32B identifies *POU5F1B* at 11,247 bp and constructs a coherent but incorrect story. *POU5F1B* is a pseudogene with weak prior evidence for a functional role in prostate cancer, while the 8q24 *MYC* super-enhancer locus is among the most-replicated cancer GWAS signals. This suggests that Qwen3-32B lacks reliable knowledge of long-range regulatory loci and defaults to a proximity heuristic.

#### Case F: Coronary Artery Disease (9p21.3 locus)

##### Context

Variant chr9_22098575_A_G; trait: coronary artery disease; gold standard gene: *CDKN2B* (p15^INK4B^). The 9p21.3 locus is the strongest common variant signal for coronary artery disease; *CDKN2B* regulates vascular smooth muscle cell proliferation. We note that a different variant at the same locus has *CDKN2A* as the gold standard gene. BioBLIP consistently predicts *CDKN2B*; Qwen3-32B consistently predicts *CDKN2A*. Each model is correct for one variant and incorrect for the other, revealing systematic paralog preference rather than case-by-case reasoning.

###### BioBLIP (*CDKN2B* ✓)

*Step 1 — Variant Annotation*. Variant type: noncoding; consequence: intron variant; nearest genes: *CDKN2B, CDKN2A, MTAP*; cell type: vascular smooth muscle cell.

*Step 2 — Gene-Level Effect*. The variant is located in the intronic region of *CDKN2B*, which encodes cyclin-dependent kinase inhibitor 2B (p15^INK4B^), a key regulator of the cell cycle. Intronic variants can affect splicing efficiency or alter regulatory elements that control gene expression. Direction: likely decreased expression or altered splicing of *CDKN2B*.

*Step 3 — Cellular Process*. Impacted pathway: cell cycle regulation, particularly the G1/S transition, via the p15^INK4B^–p16^INK4A^–Rb pathway. Cellular consequence: reduced *CDKN2B* activity may lead to increased cell proliferation and reduced cell cycle arrest in vascular smooth muscle cells, contributing to neointimal hyperplasia and atherosclerosis.

*Step 4 — Tissue-Level Effect*. Tissue: coronary arteries and vascular smooth muscle. Outcome: enhanced vascular smooth muscle cell proliferation and migration can lead to plaque formation and narrowing of coronary arteries.

*Step 5 — Disease Mechanism*. Dysregulation of *CDKN2B* in vascular smooth muscle cells promotes pathological remodelling of the arterial wall, contributing to atherosclerotic lesion development and progression of coronary artery disease.

Causal gene: *CDKN2B*.

###### Qwen3-32B (*CDKN2A ×*)

*Step 1 — Variant Consequence*. The variant 9_22098575_A_G is located on chromosome 9 at position 22,098,575. Based on its proximity to the genes *CDKN2A* and *CDKN2B* (103,274 bp and 89,270 bp away, respectively), it is likely a regulatory variant that influences gene expression. Variants in this genomic region have been associated with altered expression of *CDKN2A* and *CDKN2B*. This variant is likely an enhancer or promoter variant that modulates the transcriptional activity of *CDKN2A* and *CDKN2B*.

*Step 2 — Gene-Level Effect. CDKN2A* and *CDKN2B* encode cyclin-dependent kinase inhibitors that regulate the G1 phase of the cell cycle by inhibiting CDK4/6 activity. Reduced expression of these genes leads to unchecked cell proliferation. Direction: the variant is associated with decreased expression of *CDKN2A* and *CDKN2B*, resulting in a loss of function.

*Step 3 — Cellular Process*. Impacted pathway: cell cycle regulation, particularly the G1/S transition. Cellular consequence: reduced *CDKN2A*/*CDKN2B* activity leads to increased CDK4/6 activity, promoting uncontrolled cell proliferation.

*Step 4 — Tissue-Level Effect*. Tissue: vascular smooth muscle cells and endothelial cells in coronary arteries. Outcome: proliferation of vascular smooth muscle cells and endothelial dysfunction contribute to the formation of atherosclerotic plaques.

*Step 5 — Disease Mechanism*. The loss of *CDKN2A*/*CDKN2B* function in vascular tissues leads to increased cell proliferation and impaired cell cycle control, promoting atherosclerosis and coronary artery disease.

Causal gene: *CDKN2A*.

##### Analysis

This is the subtlest failure mode. Qwen3-32B correctly identifies the 9p21.3 locus and names both *CDKN2A* and *CDKN2B* as relevant candidates. Its biological reasoning—cell cycle dysregulation in vascular smooth muscle cells—is largely correct. However, when forced to choose a single causal gene, it defaults to *CDKN2A*, likely because *CDKN2A* (p16^INK4A^/ARF) has broader name recognition from cancer biology. BioBLIP’s explicit annotation of the variant as intronic to *CDKN2B* and the cell type as vascular smooth muscle cells resolves the ambiguity. As noted above, a mirror case at the same locus with gold gene *CDKN2A* produces the opposite outcome, indicating that each model has a fixed paralog preference at this locus rather than performing variant-specific disambiguation.

### D.5 Counterexample: Qwen3-32B Correct, BioBLIP Incorrect

Of the 26 cases where the two models disagree, BioBLIP is correct in 23 and Qwen3-32B in 3. We present the most instructive of the three counterexamples: the *HERC2*/*OCA2* pigmentation locus, where both models understand the same regulatory mechanism but differ on which gene to name as causal.

#### Case G: Skin Pigmentation — OCA2 vs. HERC2

##### Context

Variant chr15_28289631_C_T; trait: skin pigmentation; gold standard gene: *OCA2* (P protein), a melanosomal membrane transporter whose loss of expression causes oculocutaneous albinism type 2. The variant lies within intron 86 of *HERC2*, which contains a well-characterised enhancer element that controls *OCA2* expression in melanocytes.

###### Qwen3-32B (*OCA2* ✓)

*Step 1 — Variant Consequence*. The variant 15_28289631_C_T is located on chromosome 15 at position 28,289,631. It is in close proximity to *HERC2*, which is overlapping with the variant, and is also near *OCA2*, a well-known gene involved in pigmentation. This variant is likely a regulatory variant located in a promoter or enhancer region of *HERC2*, as it is in a non-coding region and is known from prior studies to influence the expression of *OCA2* through *HERC2*-mediated regulation.

*Step 2 — Gene-Level Effect. HERC2* is a regulatory gene that controls the expression of *OCA2*, which encodes a transmembrane protein involved in melanin synthesis. The variant disrupts a regulatory element that normally enhances *OCA2* expression. Direction: reduced expression of *OCA2* due to impaired regulatory function of *HERC2*.

*Step 3 — Cellular Process*. Impacted pathway: melanin biosynthesis. Cellular consequence: decreased expression of *OCA2* leads to reduced melanin production in melanocytes.

*Step 4 — Tissue-Level Effect*. Tissue: skin (melanocytes). Outcome: reduced melanin synthesis in melanocytes results in lighter skin pigmentation.

*Step 5 — Disease Mechanism*. The reduced expression of *OCA2* due to the regulatory variant in *HERC2* leads to decreased melanin production in melanocytes, which manifests as lighter skin pigmentation. This is a well-established mechanism for the genetic basis of human skin colour variation.

Causal gene: *OCA2*.

###### BioBLIP (*HERC2 ×*)

*Step 1 — Variant Annotation*. Variant type: noncoding; consequence: intron variant; nearest genes: *HERC2, OCA2*; cell type: melanocytes.

*Step 2 — Gene-Level Effect*. The variant is located in the intronic region of *HERC2*, which is a regulatory gene that influences the expression of *OCA2*, a key gene in melanin synthesis. This intronic variant likely affects the splicing or transcriptional regulation of *HERC2*, which in turn modulates *OCA2* expression. Direction: decreased *OCA2* expression due to reduced regulatory activity of *HERC2*.

*Step 3 — Cellular Process*. Impacted pathway: melanin biosynthesis. Cellular consequence: reduced melanin production in melanocytes due to lower *OCA2* activity, which is essential for the transport of tyrosine and the function of tyrosinase in melanin synthesis.

*Step 4 — Tissue-Level Effect*. Tissue: skin (melanocytes in the epidermis). Outcome: decreased melanin content in the skin, leading to lighter skin pigmentation. *Step 5 — Disease Mechanism*. The reduced melanin synthesis in melanocytes due to impaired *OCA2* expression results in decreased skin pigmentation.

Causal gene: *HERC2*.

##### Analysis

This case is notable because both models produce nearly identical biological explanations: a *HERC2* intronic variant disrupts an enhancer, reducing *OCA2* expression in melanocytes and decreasing melanin production. The two traces agree on every mechanistic step. They diverge only at the final attribution. BioBLIP names *HERC2*—the gene physically containing the variant—following its default annotation rule. Qwen3-32B names *OCA2*—the gene whose altered expression directly causes the phenotypic change—explicitly invoking “prior studies” to override the proximity anchor. For the purposes of causal gene attribution in GWAS, where the goal is to identify the gene whose dysregulation mediates the trait, *OCA2* is the correct answer. This case illustrates a limitation of BioBLIP’s proximity-based annotation: when a variant resides in a regulatory element embedded within one gene but functionally controls a neighbouring gene, BioBLIP’s default rule of naming the variant-containing gene can produce the wrong attribution, even when the downstream reasoning correctly identifies the effector gene. Qwen3-32B’s advantage here is not superior biological reasoning but memorised locus-specific knowledge sufficient to distinguish the regulatory container from the regulatory target.

## E Variant Prioritization Results

### E.1 Illustrative Reasoning Traces

#### Trace 1 Trait: HOLOPROSENCEPHALY 5 Match group: 3_prime_UTR_variant_14

Result: 13_99986571_T_C ✓ Rank 1

##### BioBLIP (*ZIC2* ✓)

Okay, let’s tackle this problem. The goal is to determine which of the 10 candidate variants is the most likely causal variant for Holoprosencephaly 5 (HPE5). All variants are intronic and noncoding, so they don’t directly affect the protein sequence. Therefore, I need to look at the nearest genes and their known associations with HPE5.

First, I recall that Holoprosencephaly is a congenital disorder involving the brain and face, and it’s often caused by mutations in genes involved in Sonic Hedgehog (SHH) signaling. HPE5 specifically is associated with mutations in the ZIC2 gene. ZIC2 is a zinc finger protein involved in neural development, and mutations here are known to cause HPE.

Looking at the list of variants, variant 9 is near ZIC2 (13_99986571_T_C). Since ZIC2 is directly linked to HPE5, this variant is a strong candidate. Even though it’s an intronic variant, it might affect gene regulation, splicing, or expression levels, which are common mechanisms for noncoding variants.

Next, I should check the other genes. Let’s go through each one:

1. NALP1: Involved in inflammation, not directly related to HPE. 2. COL4A2: Part of collagen, maybe in extracellular matrix, not directly linked. 3. RASA3: Regulates RAS signaling, but I don’t recall a direct link to HPE. 4. MAVS: Involved in antiviral response, not HPE-related. 5. GON4L: Not familiar with its role in HPE. 6. ALPPL2: Alkaline phosphatase-like, not known for HPE. 7. XGXS: Not sure about this one; might be a pseudogene or less studied. 8. MA2G4E: Part of the MA2 domain, not directly related. 9. ZIC2: As mentioned, directly associated with HPE5. 10. GPR12: A G-protein coupled receptor; no known connection to HPE.

So, variant 9 (ZIC2) is the top candidate. The others don’t have a clear link to HPE5. Even though all are intronic, the proximity to ZIC2 makes variant 9 the most likely causal. The rest are less likely because their nearest genes aren’t known to be involved in HPE5. I should rank them with variant 9 first, then the others in some order based on possible indirect links, but since none are directly connected, they would follow in arbitrary order. However, maybe some genes have roles in development that could be considered, but without specific knowledge linking them to HPE, they are lower priority.

Ranked Candidates: 1. 13_99986571_T_C 2. 13_110720333_A_T 3. 13_110429711_T_C 4. 13_113978728_A_G 5. 13_25167473_C_T 6. 13_25167401_A_G 7. 13_36302566_C_T 8. 13_101181099_C_T 9. 13_26755371_A_G 10. 13_96837203_G_A

#### Trace 2 Trait: BETA-THALASSEMIA Match group: 3_prime_UTR_variant_11

Result: 11_5225490_T_G ✓ Rank 1

##### BioBLIP (*HBB* ✓)

Okay, let’s tackle this problem. So, I need to figure out which of these 10 variants is the causal one for Beta-Thalassemia. First, I remember that Beta-Thalassemia is a genetic disorder related to the production of hemoglobin. Specifically, it’s caused by mutations in the beta-globin gene (HBB). So, the key here is to look for variants near the HBB gene.

Looking at the list of candidate variants, each has a chromosome (chr11) and nearest genes. Let’s go through them one by one.

1. Variant 11_4641757_G_A is near OR10D4, OR10D1, OR10D12. These are olfactory receptor genes, which probably aren’t related to hemoglobin. So probably not this one.
2. 11_5225490_T_G is near HBB. That’s the beta-globin gene! Since Beta-Thalassemia is caused by mutations in HBB, this variant is a strong candidate. Even though it’s an intron variant (noncoding), regulatory regions in introns can affect gene expression. So this is a top contender.
3. 11_6681654_C_G near MRPL17. MRPL17 is a mitochondrial ribosomal protein, not directly related to hemoglobin. Unlikely.
4. 11_68029165_T_C near ALDH3B1. ALDH is alcohol dehydrogenase, again not related to hemoglobin. Probably not.
5. 11_75569433_T_A near HECTD4. HECTD4 is an E3 ubiquitin ligase. Not directly related to hemoglobin. Unlikely.
6. 11_31511508_A_G near SEL1L. SEL1L is involved in ER-associated degradation. Not directly related to hemoglobin. Probably not.
7. 11_295343_G_A near GPX3. GPX3 is glutathione peroxidase, an antioxidant enzyme. Not related to hemoglobin. Unlikely.
8. 11_4641771_C_T again near OR10D4, OR10D1, OR10D12. Same as variant 1. So same reasoning, not likely.
9. 11_5268440_A_G near HBD, HBB, HBA2. HBD is delta-globin, HBB is beta-globin, and HBA2 is alpha-2 globin. Since HBB is the main gene involved in Beta-Thalassemia, this variant is also near HBB. This could be a candidate. The presence of HBB here makes it a strong possibility. However, the variant is noncoding, so maybe it’s in a regulatory region affecting HBB expression.
10. 11_74491801_T_C near ALAS2. ALAS2 is aminolevulinate synthase 2, involved in heme synthesis. Not directly related to hemoglobin’s beta chain. Unlikely.

Now, between variants 2 and 9. Both are near HBB. Let’s think about their positions. Variant 2 is at position 5,225,490, and variant 9 is at 5,268,440. The HBB gene is on chr11, and the exact location of the gene might matter. However, since both are intron variants in HBB or nearby, they could both be regulatory. But which one is more likely?

I recall that common mutations in HBB for Beta-Thalassemia include point mutations in the promoter or splice sites, but also in introns. For example, the IVS-II-654 (G>A) mutation in intron 2 of HBB is a known cause of Beta-Thalassemia. If variant 2 is in intron 2, it could be that. Alternatively, variant 9 is near HBD and HBB. However, HBD is part of the delta-globin gene cluster, which is close to HBB. So maybe variant 9 is in a region that affects HBB expression as well.

But since variant 2 is directly listed as nearest to HBB, and variant 9 is listed as nearest to HBD, HBB, HBA2. Wait, the nearest_genes for variant 9 includes HBB, so it’s in the vicinity of HBB. However, the exact position might matter. Let’s check the positions. The HBB gene is located around chr11:5,210,000 to 5,220,000 (approximate, but exact coordinates can vary). Variant 2 is at 5,225,490, which is just beyond the HBB gene. Variant 9 is at 5,268,440, which is further away. Wait, maybe I have the positions wrong. Let me think again. If HBB is around 5,210,000-5,220,000, then variant 2 is at 5,225,490, which is just after the gene. Variant 9 is at 5,268,440, which is 48 kb away. So variant 2 is closer to HBB. Therefore, variant 2 is more likely to be in the HBB gene’s regulatory region or intron, making it a better candidate.

Alternatively, maybe variant 9 is in the HBB gene itself. Wait, the consequence is “intron variant”, so if variant 9 is in an intron of HBB, then it’s a strong candidate. But the nearest_genes for variant 9 includes HBD, HBB, HBA2. So perhaps it’s in the HBB gene’s intron. However, the exact position would determine that. If variant 9 is in the HBB gene’s intron, then it’s a candidate. But the problem is that the data given is that the variant is a noncoding intron variant. So both variants 2 and 9 are in introns of HBB. But which one is more likely?

I think the key is that variant 2 is directly listed as nearest to HBB, while variant 9 is listed as nearest to HBD, HBB, HBA2. So perhaps variant 2 is within the HBB gene’s intron, making it more directly involved. Also, known mutations in HBB’s introns are known causes. For example, the IVS-II-654 (G>A) is a common mutation. If variant 2 is in a similar position, it could be the causal variant.

Therefore, variant 2 (11_5225490_T_G) is the most likely. Then variant 9 (11_5268440_A_G) is next, as it’s also near HBB. The rest are in other genes not related to hemoglobin. So the ranking would be variant 2 first, then 9, then the others in order of proximity to HBB or not. The rest are in genes unrelated to hemoglobin, so they are less likely.

Ranked Candidates: 1. 11_5225490_T_G 2. 11_5268440_A_G 3. 11_4641757_G_A 4. 11_4641771_C_T 5. 11_6681654_C_G 6. 11_68029165_T_C 7. 11_75569433_T_A 8. 11_31511508_A_G 9. 11_295343_G_A 10. 11_74491801_T_C

#### Trace 3

Trait: Immunodysregulation, Polyendocrinopathy, AND Enteropathy, X-LINKED (IPEX) Match group: `3_prime_UTR_variant_27`

Result: × Rank 8 (causal gene *FOXP3* absent from candidates)

##### BioBLIP (*FOXP3* ×)

X-linked immunodeficiency with polyendocrinopathy and enteropathy (IPEX) is a rare genetic disorder caused by mutations in the FOXP3 gene, which is located on the X chromosome. FOXP3 is a transcription factor critical for the development and function of regulatory T cells (Tregs), which are essential for maintaining immune tolerance and preventing autoimmune diseases. However, none of the candidate variants listed are in or near the FOXP3 gene. This suggests that either the variants are in linkage disequilibrium with the true causal variant in FOXP3, or they are in other genes that may play a role in the same biological pathways or processes related to IPEX.

Given that all variants are intronic and noncoding, their potential impact on gene function would be through regulatory mechanisms, such as affecting splicing, transcription factor binding, or enhancer activity. The likelihood of a variant being causal can be inferred based on the biological relevance of the nearest genes to the immune system, endocrine function, and intestinal health.

Let’s analyze the nearest genes for each variant:

1. **X_136214250_C_T (MPAN)**: MPAN is a poorly characterized gene with no clear link to immune regulation, endocrine function, or intestinal health. 2. **X_41696942_A_T (GPX7, GPR146)**: GPX7 is a glutathione peroxidase involved in antioxidant defense, which could be relevant to immune function. GPR146 is a G protein-coupled receptor with unknown function in the context of IPEX. 3. **X_20125252_G_T (EIF1AX, EIF1A, EIF1AY)**: EIF1A and EIF1AY are eukaryotic translation initiation factors. While they are essential for protein synthesis, their direct role in immune regulation, endocrine function, or intestinal health is not well established. 4. **X_49250456_T_C (COPZ2)**: COPZ2 is involved in COPI-coated vesicle formation and intracellular trafficking. It has no clear link to IPEX. 5. **X_111689561_G_C (ALG13)**: ALG13 is an asparagine-linked glycosylation enzyme. Glycosylation is important for protein function, but its role in IPEX is not clear. 6. **X_48608798_C_T (WDR72)**: WDR72 is a WD repeat-containing protein with no clear link to IPEX. 7. **X_155057576_C_G (DCUN1D2, TMEM132D4, TMEM132D3)**: These genes are involved in endosomal trafficking and membrane dynamics, but their relevance to IPEX is unclear. 8. **X_143626437_T_C (SLITRK4)**: SLITRK4 is a receptor for SLIT proteins, which are involved in axon guidance and cell migration. Its role in immune regulation is not well established. 9. **X_145825876_A_G (SLC25A10)**: SLC25A10 is a mitochondrial carrier protein involved in the transport of aspartate and glutamate. It has no clear link to IPEX. 10. **X_111519499_T_C (STIMATE2)**: STIMATE2 is a poorly characterized gene with no clear link to IPEX.

Given the lack of clear links to IPEX-related pathways, it is possible that the true causal variant is in linkage disequilibrium with one of these variants, particularly if it is in or near the FOXP3 gene. However, since none of the listed variants are in or near FOXP3, the most likely causal variant would be the one with the nearest gene that has the strongest potential link to immune regulation, endocrine function, or intestinal health.

Based on this analysis, the most likely causal variant is **X_41696942_A_T (GPX7, GPR146)**, as GPX7 is involved in antioxidant defense, which could be relevant to immune function. The next most likely candidates would be those with genes involved in more general cellular processes, such as **X_20125252_G_T (EIF1AX, EIF1A, EIF1AY)** and **X_48608798_C_T (WDR72)**, as they are involved in protein synthesis and cellular processes that could indirectly affect immune function.

Ranked Candidates: 1. X_41696942_A_T 2. X_20125252_G_T 3. X_48608798_C_T 4. X_155057576_C_G 5. X_143626437_T_C 6. X_145825876_A_G 7. X_111689561_G_C 8. X_49250456_T_C 9. X_111519499_T_C 10. X_136214250_C_T

## References

[1] Eric Wang, Samuel Schmidgall, Paul F Jaeger, Fan Zhang, Rory Pilgrim, Yossi Matias, Joelle Barral, David Fleet, and Shekoofeh Azizi. Txgemma: Efficient and agentic llms for therapeutics. arXiv preprint 2504.06196, 2025.

[2] Qizhi Pei, Wei Zhang, Jinhua Zhu, Kehan Wu, Kaiyuan Gao, Lijun Wu, Yingce Xia, and Rui Yan. Biot5: Enriching cross-modal integration in biology with chemical knowledge and natural language associations. In Proceedings of the 2023 Conference on Empirical Methods in Natural Language Processing, pages 1102–1123, 2023.

[3] Ross Taylor, Marcin Kardas, Guillem Cucurull, Thomas Scialom, Anthony Hartshorn, Elvis Saravia, Andrew Poulton, Viktor Kerkez, and Robert Stojnic. Galactica: A large language model for science. arXiv preprint 2211.09085, 2022.

[4] Žiga Avsec, Natasha Latysheva, Jun Cheng, Guido Novati, Kyle R Taylor, Tom Ward, Clare Bycroft, et al. Advancing regulatory variant effect prediction with alphagenome. Nature, 649(8099):1206–1218, 2026.

[5] Josh Abramson, Jonas Adler, Jack Dunger, Richard Evans, Tim Green, Alexander Pritzel, Olaf Ronneberger, Lindsay Willmore, Andrew J Ballard, Joshua Bambrick, et al. Accurate structure prediction of biomolecular interactions with alphafold 3. Nature, 630(8016):493–500, 2024.

[6] Garyk Brixi, Matthew G Durrant, Jerome Ku, Mohsen Naghipourfar, Michael Poli, Gwanggyu Sun, Greg Brockman, Daniel Chang, Alison Fanton, Gabriel A Gonzalez, et al. Genome modelling and design across all domains of life with evo 2. Nature, pages 1–13, 2026.

[7] Adibvafa Fallahpour, Andrew Magnuson, Purav Gupta, Shihao Ma, Jack Naimer, Arnav Shah, Haonan Duan, Omar Ibrahim, Hani Goodarzi, Chris J Maddison, et al. Bioreason: Incentivizing multimodal biological reasoning within a dna-llm model. arXiv preprint 2505.23579, 2025.

[8] Adibvafa Fallahpour, Arman Seyed-Ahmadi, Parsa Idehpour, Omar Ibrahim, Purav Gupta, Jack Naimer, Kevin Zhu, Arnav Shah, Shihao Ma, Abhinav Adduri, Talu Güloglu, Nuo Liu, Haotian Cui, Arihant Jain, Max de Castro, Amirfaham Fallahpour, Antonio Cembellin-Prieto, John S. Stiles, Filip Nemčko, Alexander A. Nevue, Hyungseok C. Moon, Lucas Sosnick, Olivia Markham, Haonan Duan, Michelle Y. Y. Lee, Andrea F. M. Salvador, Chris J. Maddison, Christoph A. Thaiss, Chiara Ricci-Tam, Brian S. Plosky, Dave P. Burke, Patrick D. Hsu, Hani Goodarzi, and Bo Wang. Bioreason-pro: Advancing protein function prediction with multimodal biological reasoning. bioRxiv, 2026.

[9] Yijia Xiao, Edward Sun, Yiqiao Jin, Qifan Wang, and Wei Wang. Proteingpt: Multimodal llm for protein property prediction and structure understanding. arXiv preprint 2408.11363, 2024.

[10] Hadi Abdine, Michail Chatzianastasis, Costas Bouyioukos, and Michalis Vazirgiannis. Prot2text: Multimodal protein’s function generation with gnns and transformers. In Proceedings of the AAAI conference on artificial intelligence, volume 38, pages 10757–10765, 2024.

[11] Ching-Huei Tsou, Michal Ozery-Flato, Ella Barkan, Diwakar Mahajan, and Ben Shapira. Bioverse: Representation alignment of biomedical modalities to llms for multi-modal reasoning. arXiv preprint 2510.01428, 2025.

[12] Yoel Shoshan, Moshiko Raboh, Michal Ozery-Flato, Vadim Ratner, Alex Golts, Jeffrey K Weber, Ella Barkan, Simona Rabinovici-Cohen, Sagi Polaczek, Ido Amos, et al. Mammal–molecular aligned multi-modal architecture and language. arXiv preprint 2410.22367, 2024.

[13] Bernardo P de Almeida, Guillaume Richard, Hugo Dalla-Torre, Christopher Blum, Lorenz Hexemer, Priyanka Pandey, Stefan Laurent, Chandana Rajesh, Marie Lopez, Alexandre Laterre, et al. A multimodal conversational agent for dna, rna and protein tasks. Nature Machine Intelligence, 7(6):928–941, 2025.

[14] Gonzalo Benegas, Gökcen Eraslan, and Yun S Song. Benchmarking dna sequence models for causal regulatory variant prediction in human genetics. bioRxiv, 2025.

[15] Céline Marquet, Michael Heinzinger, Tobias Olenyi, Christian Dallago, Kyra Erckert, Michael Bernhofer, Dmitrii Nechaev, and Burkhard Rost. Embeddings from protein language models predict conservation and variant effects. Human genetics, 141(10):1629–1647, 2022.

[16] Shashank Yadav, Dhvani Sandip Vora, Durai Sundar, and Jaspreet Kaur Dhanjal. Tcr-esm: Employing protein language embeddings to predict tcr-peptide-mhc binding. Computational and Structural Biotechnology Journal, 23:165–173, 2024.

[17] Junnan Li, Dongxu Li, Silvio Savarese, and Steven Hoi. Blip-2: Bootstrapping language-image pre-training with frozen image encoders and large language models. In International conference on machine learning, pages 19730–19742. PMLR, 2023.

[18] An Yang, Anfeng Li, Baosong Yang, Beichen Zhang, Binyuan Hui, Bo Zheng, Bowen Yu, Chang Gao, Chengen Huang, Chenxu Lv, et al. Qwen3 technical report. arXiv preprint 2505.09388, 2025.

[19] Wenliang Dai, Junnan Li, Dongxu Li, Anthony Tiong, Junqi Zhao, Weisheng Wang, Boyang Li, Pascale N Fung, and Steven Hoi. Instructblip: Towards general-purpose vision-language models with instruction tuning. Advances in neural information processing systems, 36:49250–49267, 2023.

[20] Yiqun Chen and James Zou. Genept: a simple but effective foundation model for genes and cells built from chatgpt. BioRxiv, pages 2023–10, 2024.

[21] Zeming Lin, Halil Akin, Roshan Rao, Brian Hie, Zhongkai Zhu, Wenting Lu, Nikita Smetanin, Robert Verkuil, Ori Kabeli, Yaniv Shmueli, et al. Evolutionary-scale prediction of atomic-level protein structure with a language model. Science, 379(6637):1123–1130, 2023.

[22] Annalisa Buniello, Daniel Suveges, Carlos Cruz-Castillo, Manuel Bernal Llinares, Helena Cornu, Irene Lopez, Kirill Tsukanov, Juan María Roldán-Romero, Chintan Mehta, Luca Fumis, Graham McNeill, James D Hayhurst, Ricardo Esteban Martinez Osorio, Ehsan Barkhordari, Javier Ferrer, Miguel Carmona, Prashant Uniyal, Maria J Falaguera, Polina Rusina, Ines Smit, Jeremy Schwartzentruber, Tobi Alegbe, Vivien W Ho, Daniel Considine, Xiangyu Ge, Szymon Szyszkowski, Yakov Tsepilov, Maya Ghoussaini, Ian Dunham, David G Hulcoop, Ellen M McDonagh, and David Ochoa. Open targets platform: facilitating therapeutic hypotheses building in drug discovery. Nucleic Acids Research, 53(D1):D1467–D1475, 01 2025.

[23] Manning Lab. Genelocator: Variant annotation by mapping chromosome positions to gene names. https://github.com, 2019.

[24] Gonzalo Benegas, Carlos Albors, Alan J Aw, Chengzhong Ye, and Yun S Song. A dna language model based on multispecies alignment predicts the effects of genome-wide variants. Nature Biotechnology, 43(12):1960–1965, 2025.

[25] Shilong Liu, Hao Cheng, Haotian Liu, Hao Zhang, Feng Li, Tianhe Ren, Xueyan Zou, Jianwei Yang, Hang Su, Jun Zhu, et al. Llava-plus: Learning to use tools for creating multimodal agents. In European conference on computer vision, pages 126–142. Springer, 2024.

[26] Qiao Jin, Yifan Yang, Qingyu Chen, and Zhiyong Lu. Genegpt: augmenting large language models with domain tools for improved access to biomedical information. Bioinformatics, 40(2):btae075, 2024.

[27] Mohammad Asadi, Jack W O’Sullivan, Fang Cao, Tahoura Nedaee, Kamyar Fardi, Fei-Fei Li, Ehsan Adeli, and Euan Ashley. Mirage the illusion of visual understanding. arXiv preprint 2603.21687, 2026.

[28] Moritz Schaefer, Peter Peneder, Daniel Malzl, Anna Hakobyan, Varun S Sharma, Thomas Krausgruber, Jörg Menche, Eleni Tomazou, and Christoph Bock. Joint embedding of transcriptomes and text enables interactive single-cell rna-seq data exploration via natural language. In ICLR 2024 Workshop on Machine Learning for Genomics Explorations, 2024.

[29] Moritz Schaefer, Peter Peneder, Daniel Malzl, Salvo Danilo Lombardo, Mihaela Peycheva, Jake Burton, Anna Hakobyan, Varun Sharma, Thomas Krausgruber, Celine Sin, et al. Multimodal learning enables chat-based exploration of single-cell data. Nature Biotechnology, pages 1–11, 2025.

[30] Yiqing Shen, Zan Chen, Michail Mamalakis, Yungeng Liu, Tianbin Li, Yanzhou Su, Junjun He, Pietro Liò, and Yu Guang Wang. Toursynbio: A multi-modal large model and agent framework to bridge text and protein sequences for protein engineering. In 2024 IEEE International Conference on Bioinformatics and Biomedicine (BIBM), pages 2382–2389. IEEE, 2024.

[31] Mingze Yin, Hanjing Zhou, Yiheng Zhu, Miao Lin, Yixuan Wu, Jialu Wu, Hongxia Xu, Chang-Yu Hsieh, Tingjun Hou, Jintai Chen, et al. Multi-modal clip-informed protein editing. Health Data Science, 4:0211, 2024.

[32] Yujia Wang, Jihong Guan, Wengen Li, Shuigeng Zhou, and Xuhong Wang. Biobridge: Bridging proteins and language for enhanced biological reasoning with llms. In 2025 IEEE International Conference on Bioinformatics and Biomedicine (BIBM), pages 70–75. IEEE, 2025.

